# Lack of mucosal cholinergic innervation is associated with increased risk of enterocolitis in Hirschsprung’s disease

**DOI:** 10.1101/2020.06.15.151621

**Authors:** Simone Keck, Virginie Galati-Fournier, Urs Kym, Michèle Moesch, Jakob Usemann, Ulrike Subotic, Sasha J. Tharakan, Thomas Krebs, Eleuthere Stathopoulos, Peter Schmittenbecher, Dietmar Cholewa, Philipp Romero, Bertram Reingruber, Elisabeth Bruder, NIG study Group, Stefan Holland-Cunz

**Affiliations:** Department of Pediatric Surgery, University Children’s Hospital Basel (UKBB) and University of Basel, Basel, Switzerland; Department of Pediatric Pulmonology, University Children’s Hospital Basel (UKBB), Basel and Division of Respiratory Medicine, University Children’s Hospital Zurich, Zurich, Switzerland; Department of Pediatric Surgery, University Children’s Hospital Zurich, Zurich, Switzerland; Department of Pediatric Surgery, Children’s Hospital of Eastern Switzerland, St. Gallen, Switzerland; Department of Pediatric Surgery, University Hospital of Lausanne (CHUV), Lausanne, Switzerland; Department of Pediatric Surgery, Municipal Hospital, Karlsruhe, Germany; Department of Pediatric Surgery, University Hospital of Bern, Bern, Switzerland; Department of Pediatric Surgery, University Hospital of Heidelberg, Heidelberg, Germany; Department of Pediatric Surgery, Florence Nightingale Hospital, Düsseldorf, Germany; Department of Medical Genetics and Pathology, University Hospital Basel, Basel, Switzerland; Department of Pathology, University Hospital of Lausanne (CHUV) and University of Lausanne, Lausanne, Switzerland

**Keywords:** Enterocolitis, Neuroimmunology, Cholinergic neurons, Macrophages, Th17 cells, Microbiome

## Abstract

Hirschsprung’s disease (HSCR) is a congenital intestinal motility disorder defined by the absence of enteric nervous cells (ganglia). The development of HSCR-associated enterocolitis remains a life-threatening complication. Absence of enteric ganglia implicates extramural innervation of acetylcholine-secreting (cholinergic) nerve fibers. Cholinergic signals have been reported to control excessive inflammation, but the impact on HSCR-associated enterocolitis is unknown.

**Methods:** We enrolled 44 HSCR patients in a prospective multicenter study and grouped them according to their degree of colonic mucosal cholinergic innervation using immunohistochemistry. The fiber phenotype was correlated with the tissue cytokine profile as well as immune cell frequencies using quantitative reverse-transcribed real-time polymerase chain reaction (qRT-PCR) of whole colonic tissue and fluorescence-activated cell sorting (FACS) analysis of isolated colonic immune cells. Fiber-associated immune cells were identified using confocal immunofluorescence microscopy and characterized by RNA-seq analysis. Microbial dysbiosis was analyzed in colonic patient tissue using 16S rDNA gene sequencing. Finally, the fiber phenotype was correlated with postoperative enterocolitis manifestation.

**Results:** We provided evidence that extrinsic mucosal innervation correlated with reduced interleukin (IL)-17 cytokine levels and T-helper-17 (Th17) cell frequencies. Bipolar CD14^high^ macrophages colocalized with neurons and expressed significantly less interleukin-23, a Th17-promoting cytokine. HSCR patients lacking mucosal cholinergic nerve fibers showed microbial dysbiosis and had a higher incidence of postoperative enterocolitis.

**Conclusion:** The mucosal fiber phenotype might serve as a new prognostic marker for enterocolitis development in HSCR patients and may offer an approach to personalized patient care and new future therapeutic options. (www.clinicaltrials.gov accessing number NCT03617640)

## Main

Hirschsprung’s disease (HSCR) is a multigenetic congenital malformation of the colon characterized by a distal lack of enteric ganglia cells (aganglionosis) leading to intestinal obstruction and prestenotic megacolon. In affected infants, the aganglionic bowel part is surgically removed, and healthy ganglionic tissue is pulled through and reconnected to the anus. Either pre- or postoperatively, 20% to 50% of patients suffer from life-threatening HSCR-associated enterocolitis leading to acute abdominal distension, diarrhea, and fever up to sepsis and organ failure^1–3^. The underlying pathogenesis is not fully understood but appears to involve changes in intestinal barrier function^4–6^, immune response^7–9^, and the microbiome^10–15^.

As a consequence of lacking an intrinsic enteric nervous system (ENS), extrinsic acetylcholine (ACh)-producing (cholinergic) nerve fibers originating from sacral roots S2-S4 innervate the distal colon. ACh released from these fibers is responsible for constant muscle contraction in HSCR patients^16^.

Intestinal cholinergic signals not only coordinate physiologic bowel function but additionally prevent excessive inflammation by allowing the intestinal immune system to tolerate harmless commensals and food antigens^17^. The cholinergic-antiinflammatory-pathway (CAIP) was first described by the Tracey group^18^. In a model of systemic inflammation, vagus nerve (VN) stimulation favored the release of ACh by memory T cells which in turn bound to α7-nicotinic ACh receptor (α7nAChR)- positive splenic macrophages (MΦ) to reduce tumor necrosis factor (TNF)-α production^18,19^. This concept was confirmed in intestinal inflammation models involving colitis^20–23^, food allergy^24^, postoperative ileus (POI)^25^, and pancreatitis using VN stimulation or pharmacologic targeting of α7nAChR. In humans, nicotine, which activates nAChRs, has been associated with clinical improvement of ulcerative colitis^26^. VN stimulation in Crohn’s disease results in amelioration of disease activity^27^. Contrary to systemic CAIP, intestinal modulation is independent of the spleen and involves direct activation of local enteric neurons^25,28,29^. Although VN activation leads to downregulation of inflammatory cytokine production by muscularis MΦ via α7AChR activation in POI, it remains unclear which receptors and cell types are involved in colitis amelioration^30,31^. Given the fact that the ENS utilizes more than 30 different neurotransmitters, it is challenging to identify individual immune-modulating functions. As HSCR patients miss these ENS-specific neurotransmitters, the study of cholinergic neurons is possible.

Here, we investigated the role of cholinergic mucosal innervation in regulating the immune cell status, microbial composition, and development of enterocolitis in HSCR patients.

## Materials and Methods

Detailed protocols are provided in the Supplementary Materials and Methods

## Results

### Distinct cholinergic innervation of the colon in HSCR patients

In this prospective, multicenter study, we collected fresh tissue of different colonic segments from 44 pediatric HSCR patients undergoing therapeutic pull-through surgery as well as 6 control patients (non-HSCR) undergoing surgery for miscellaneous reasons. The length of aganglionic colon was determined by experienced pathologists of the respective pathology units (see Methods section, Supplementary Table 1 and Supplementary Table 2)^32^. Compared to non-HSCR colonic tissue, the aganglionic colon of HSCR patients lack the myenteric and submucosal plexus but show the characteristic increase of ACh esterase (AChE) activity in the rectosigmoid (Fig. 1a). Increased cholinergic activity is limited to the aganglionic rectosigmoid distal colon and decreases in caudocranial direction^16,33^. As a consequence, no extrinsic mucosal AChE^+^ nerve fibers can be detected in the proximal aganglionic colon descendens (CD), with only few present in the intermuscular space. Depending on the length of aganglionosis, the proximal CD segment of HSCR patients can be ganglionic or aganglionic (Fig. 1a and Supplementary Table 1). CD segments showing a hypoganglionic phenotype (transition zone) were excluded from the analysis.

**Fig. 1.**
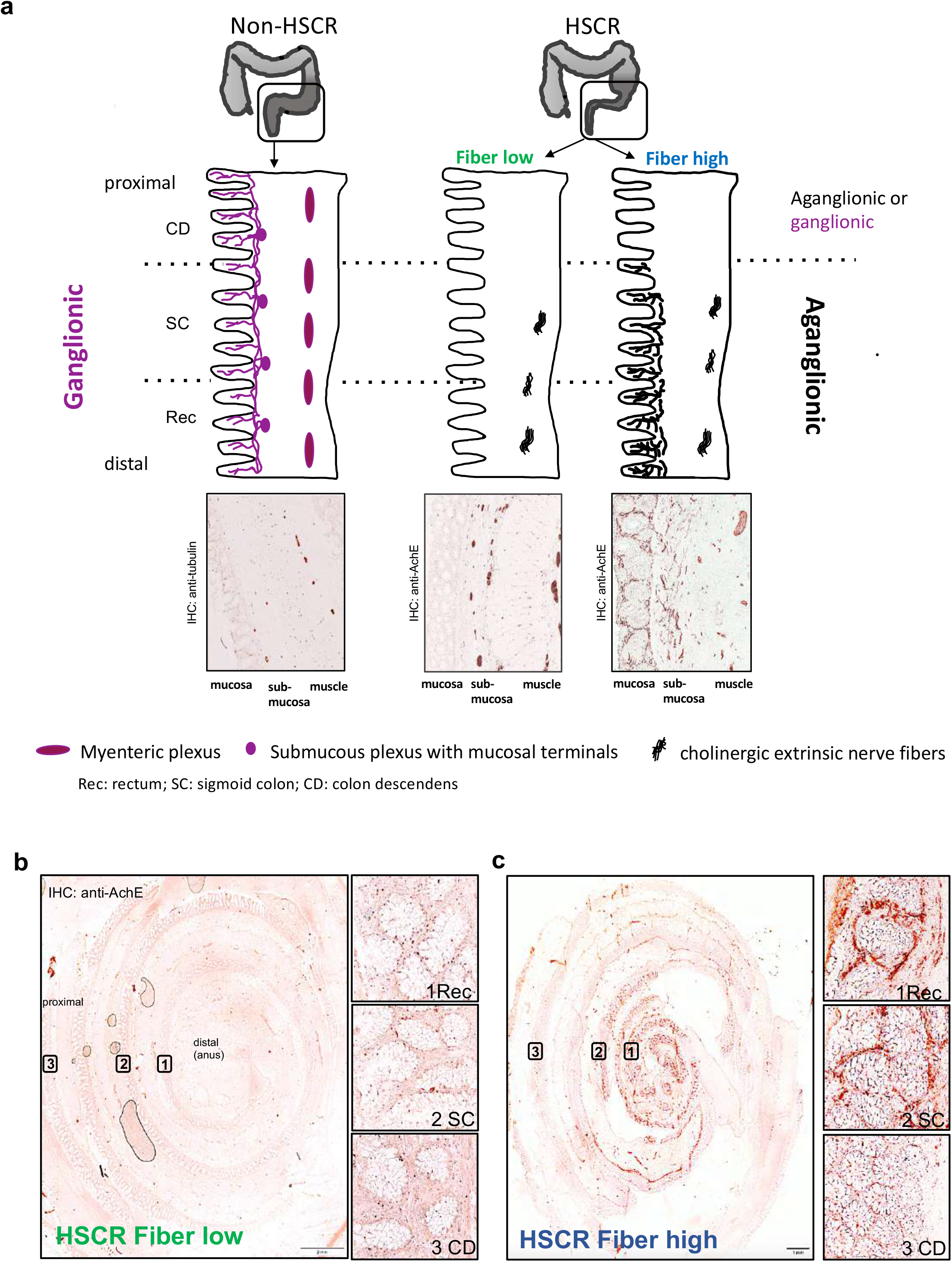
Distribution of neuronal innervation and fiber scoring of HSCR patients. **a,** Schematic representation of neuronal innervation in non-HSCR (control group) and HSCR patients. Images (5 μm colonic cryosections) show immunohistochemical tubulin staining of ganglionic CD region (non-HSCR) or AChE staining of aganglionic rectum region (HSCR). CD regions of HSCR patients were ganglionic or aganglionic dependent on the length of aganglionosis (see Supplementary Table 1). Rec (rectum); SC (sigmoid colon); CD (colon descendens). **b-c**, Longitudinal colonic strips were rolled up from the distal to proximal ends and cryopreserved as a ‘Swiss roll’. Cholinergic nerve fibers were visualized in 5 μm cryosections by AChE immunohistochemical staining using brightfield microscopy. The presence and density of AChE^+^ fibers in rectum (1 REC) and sigmoid colon (2 SC) were evaluated by four individuals. **b**, Representative image of a fiber-low colonic sections. **c**, Representative image of a fiber-high colonic section. Close-up view shows epithelial region from the different colonic segments.

We performed anti-AChE immunohistochemistry staining in cryosections of longitudinally rolled colonic stripes (Swiss roll), including rectum (REC), sigmoid colon (SC), and colon descendens (CD) from HSCR patients. All patient tissues showed the typical thick AChE^+^ fibers in the muscle region, but the density of thin mucosal fibers varied between patients (Fig. 1b, c). According to mucosal innervation in the distal rectosigmoid colon, the sections were semi-quantitatively allocated into “fiber-low” (Fig. 1b) and “fiber-high” (Fig. 1c) patient groups in a double-blind investigation (see methods for details). Fiber phenotype was validated using tubulin immunohistochemistry (Supplementary Figure 1). Consequently, for further analysis, aganglionic and ganglionic segments with variable mucosal cholinergic innervation were defined as illustrated by anti-β3tubulin (pan neuronal marker) and anti-AChE (cholinergic marker) immunofluorescent staining (Fig. 2a).

**Fig. 2.**
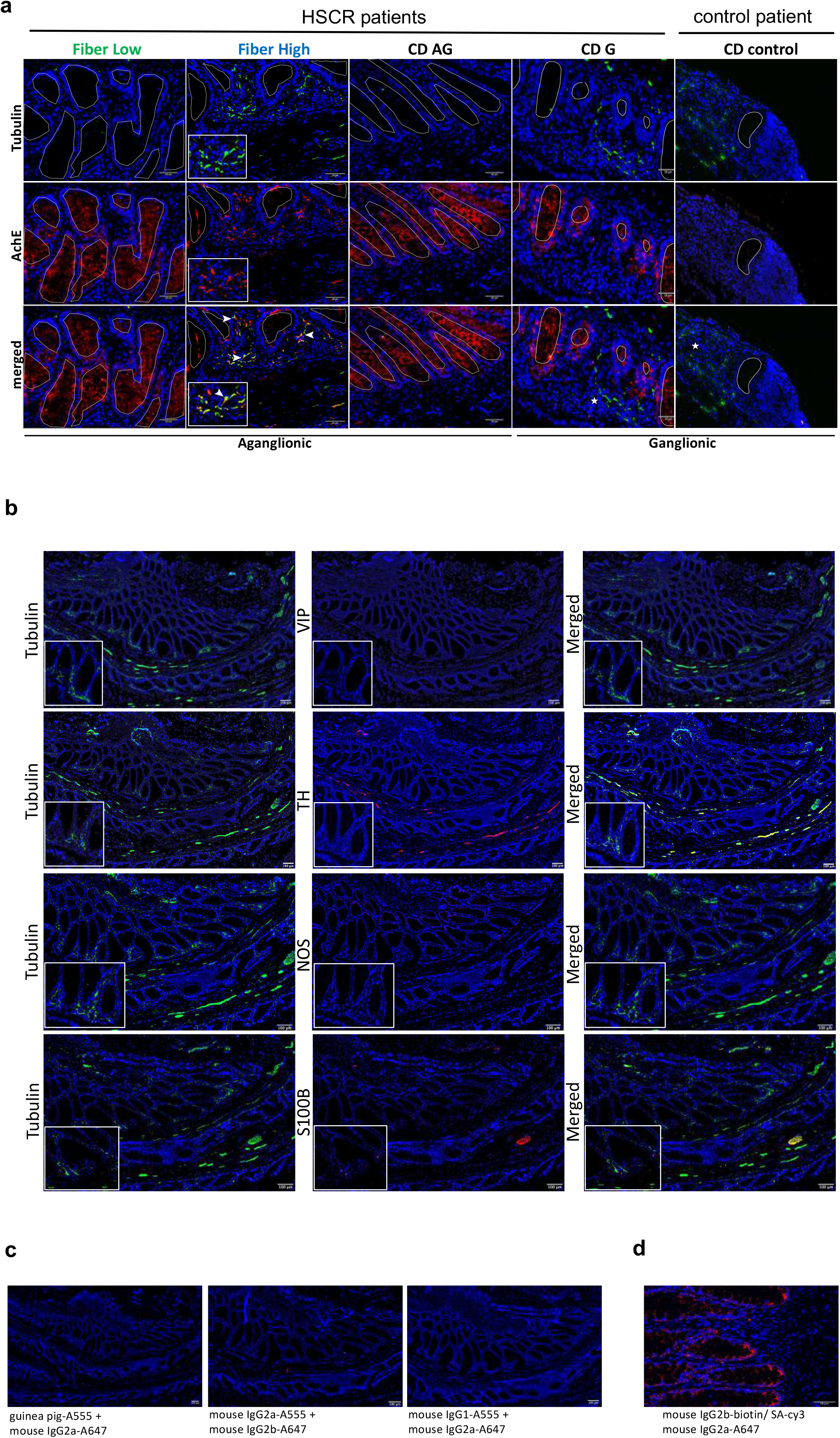
Mucosal cholinergic innervation in distal aganglionic colon of fiber high HSCR patient. **a,** Differently innervated colonic segments from aganglionic and ganglionic regions of HSCR patients and control patients used for subsequent laboratory analysis. Immunofluorescence (5 μm cryosections) of epithelial/mucosal region using tubulin (Alexa647, green), a general neuronal marker, and AChE (Cy3, red), a cholinergic marker. DAPI (blue) shows cell nuclei. Images were taken by fluorescence microscopy (20x, scale bar 50 μm) and processed by Fiji software. Encircled areas mark the lumen. Arrows mark cholinergic fibers; asterisk mark non-cholinergic fibers. **b**, Immunofluorescence (5 μm Swiss roll cryosections) of aganglionic rectum region from a fiber-high patient using tubulin (Alexa647, green), a general neuronal marker, combined with VIP (vasoactive intestinal peptide; A555, red), labeling peptidergic neurons; TH (tyrosine hydroxylase; A555, red), labeling dopaminergic and adrenergic neurons; NOS (nitric oxide synthase, A555, red), labeling nitrinergic neurons and S100B (S100 calcium-binding protein B, A555, red), labeling glia cells. DAPI (blue) shows cell nuclei. Images were taken by fluorescence microscopy (20x, scale bar 50 μm) and processed by Fiji software. Close-up views (white boxes) show mucosal/epithelial region. **c-d**, Immunofluorescence (5 μm Swiss roll cryosections) of aganglionic rectum region from a fiber-high patient using secondary antibody controls.

Thus, rectosigmoid fiber-low tissues were those lacking intrinsic nerve cell bodies as well as extrinsic mucosal cholinergic innervation. Rectosigmoid Fiber-high tissues were those lacking intrinsic nerve cell bodies but showing extrinsic mucosal cholinergic innervation (Fig. 2a, arrow). Aganglionic CD sections from HSCR patients were defined as those lacking intrinsic nerve cell bodies as well as extrinsic mucosal cholinergic innervation (because of extrinsic nerve fiber restriction to the rectosigmoid). Hence, aganglionic CD segments were used from both fiber-low and fiber-high patient tissue. Ganglionic CD sections from HSCR and non-HSCR patients showed the presence of non-cholinergic intrinsic nerve cell bodies (ganglia) with mucosal nerve terminals (Fig. 2a, asterisk) and were considered as normally innervated. Secondary antibody controls showed a high background in the epithelial crypt region for AChE due to biotin/streptavidin amplification (Fig. 2d). By performing immunofluorescence studies in fiber-high tissue, we could exclude peptidergic (VIP), dopaminergic/adrenergic (TH), and nitrinergic (NOS) origins of the mucosal fibers (Fig. 2b). However, we detected TH^+^ nerve bundles in the submucosa and muscle region. Sporadic fiber bundles were positive for the glia cell marker S100B, and the majority of mucosal fibers were not associated with glia cells (Fig. 2b). Respective secondary antibody controls were negative (Fig. 2c). We did not observe any significant patient bias in terms of anthropometric, clinical, environmental, or maternal factors between fiber-low and fiber-high tissues, except a higher percentage of infants delivered by cesarean section in the fiber-high patient group (Supplementary Table 3).

### Absence of cholinergic fibers correlated with inflammatory immune status

A functional ENS is crucial for intestinal immune homeostasis^34^. To investigate if the absence of neuronal signals impairs the overall inflammatory cytokine status in HSCR patients, we performed quantitative real-time polymerase chain reaction (q-RT PCR) analysis of whole colonic tissue from HSCR patients and control patients. TNF-α did not significantly vary between the different colonic segments from HSCR patients and control patients (Fig. 3a, Supplementary Table 4). Cytokines involved in Rorγt^+^ Th17 cell differentiation, i.e., IL-1β, IL-23, and IL-6^35,36^, were significantly elevated in aganglionic CD segments compared to ganglionic CD segments, but they were not significantly changed compared to fiber-high segments. IL-17 and IL-22, two Th17 effector cytokines, were significantly elevated in aganglionic CD sections compared to ganglionic CD sections from HSCR patients (Fig. 3a, Supplementary Table 4). Both cytokines were significantly attenuated in cholinergically innervated fiber-high segments compared to fiber-low or aganglionic CD segments.

**Fig. 3.**
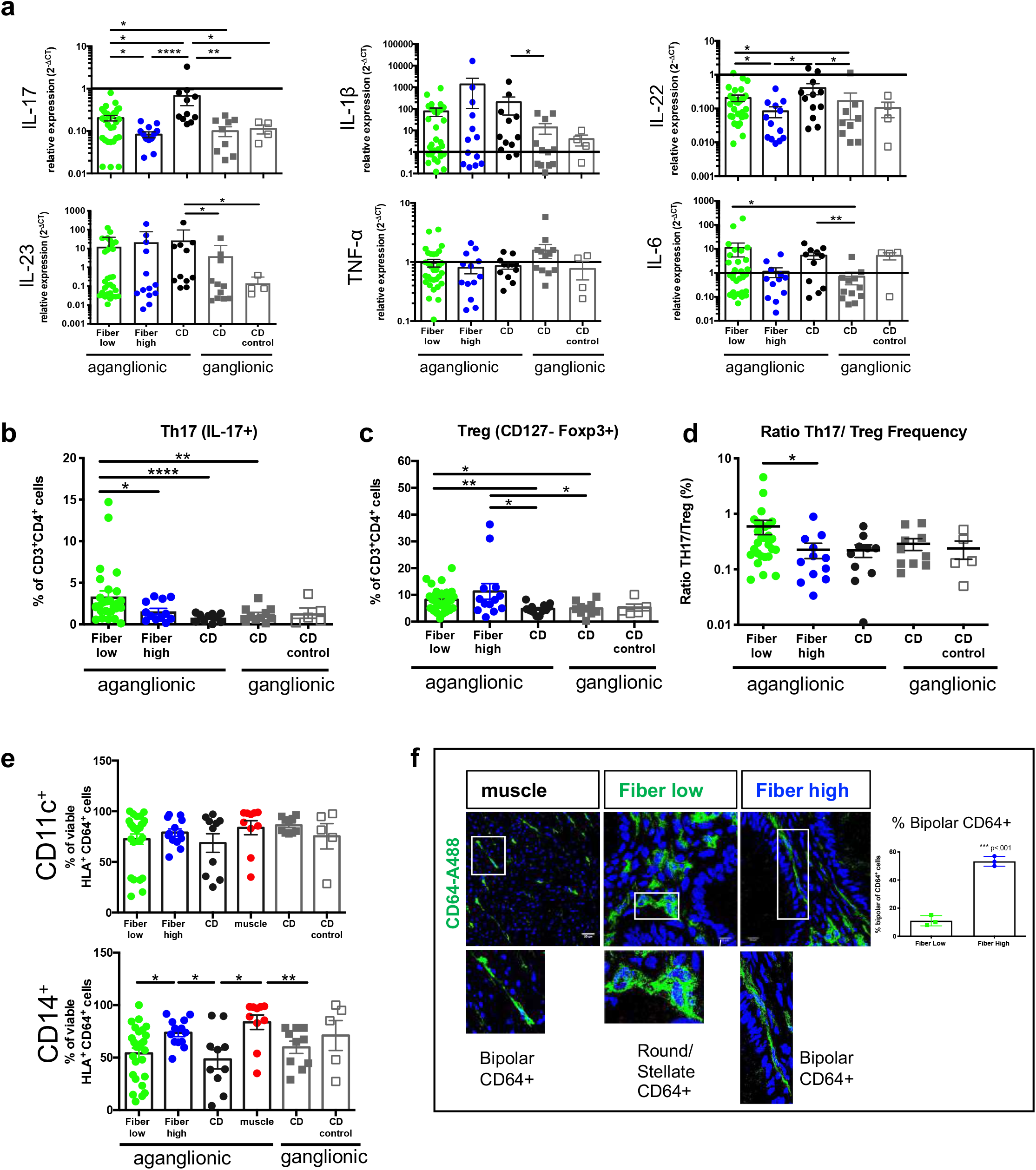
Cytokine profile and immune cell populations in colonic tissue. **a,** Quantitative RT-PCR analysis of total colonic tissue isolated from different aganglionic (AG) and ganglionic (G) colonic segments of HSCR and control patients. Fiber-low (n=31); fiber-high (n=13); AG CD (n=12); G CD (n=11); control CD (n=4). IL-17 expression was not detectable in 1 AG CD sample and in 1 G CD sample. IL-22 expression was not detectable in 2 G CD sample. **b-d**, Flow cytometric analysis of mononuclear cells isolated from different aganglionic and ganglionic colonic segments of HSCR and control patients. **b**, Frequencies of IL-17^+^ Th17 T cells were determined in viable CD3^+^CD4^+^ lymphocytes restimulated with PMA/Ionomycin (Supplementary Fig. 3a). **c**, Treg cells were defined as CD127^-^Foxp3^+^ cells in unstimulated viable CD3^+^CD4^+^ lymphocytes using Foxp3/transcription factor staining buffer set (Supplementary Fig. 3b). Fiber-low (n=30); fiber-high (n=13); AG CD (n=10); G CD (n=10); control CD (n=5) **d,** Th17/Treg ratio is shown. **e**, Frequencies of CD11c^+^ and CD14^+^ MΦ were evaluated in viable HLA^+^CD64^+^ cells by flow cytometry (Supplementary Fig. 2d). Fiber-low (n=26); Fiber-high (n=13); AG CD (n=10); muscle (n=10); G CD (n=10); control CD (n=5). **f,** CD64^+^ MΦ were assessed by immunofluorescence in 5 μm colonic cryosections from a muscle region (left), a mucosal region from a fiber-low patient, and a mucosal region from fiber-high patient (right). Scale bar 30 μm (muscle) and 10 μm (fiber-low; fiber-high). Images from 3 fiber-high and 3 fiber-low patients were analyzed for MΦ shape. A total of 100-200 CD64^+^ cells per patient were counted and the percentage of bipolar cells calculated (bar graph). Scatter plots show means ± SEM. Significance was determined using unpaired nonparametric Mann-Whitney test (* p≤ 0.05; ** p≤ 0.01; *** p≤ 0.001). Source data are provided as a Source Data file.

Beside innate lymphoid cells 3 (ILC3) and others, Th17 cells are the main producers of IL-17 and IL-22 and are counter-regulated by forkhead box P3 positive (Foxp3^+^) regulatory T cells (Treg)^37–40^. Frequencies of mucosal IL17^+^ Th17 and Foxp3^+^ Treg T cells were assessed by flow cytometry (Fig. 3 b, c, supplementary Fig. 3a, b). Th17 frequencies were significantly increased in fiber-low segments compared to fiber-high segments or CD segments (Fig. 3b, Supplementary Fig. 2a showing the percentage of total cells). We observed no difference between aganglionic and ganglionic CD tissue of HSCR and control patients. Treg frequencies were significantly elevated in both fiber-low and fiber-high tissues from HSCR patients (Fig. 3c, Supplementary Fig. 2a showing the percentage of total cells). However, the ratio of Th17/Treg was significantly higher in fiber-low segments compared to fiber-high segments (Fig. 3d, Supplementary Fig. 2a showing the percentage of total cells). Frequencies of natural killer (NK) cell receptors (NCR)^+^ and NCR^-^ ILC3s were comparable between fiber-low and fiber-high tissues from HSCR patients. However, the NCR^-^ ILC3 population was significantly elevated in fiber-low colonic segments compared to ganglionic CD of HSCR patients (Supplementary Fig. 2b). Thus, lack of an ENS appeared to have an impact on inflammatory cytokine production, whereas lack of cholinergic neurons seemed to affect Th17 cells and their related cytokines. CX3CR1^+^ MΦ represent the major antigen-presenting cell (APC) type in the colon and are crucial for CD4 T cell differentiation^41^. These MΦ exhibit a non-inflammatory M2-like phenotype and are essential for local expansion and maintenance of Treg^42,14^. In contrast, intestinal CCR2^+^ CX3CR1^+^ blood-monocyte-derived MΦ are essential for Th17 cell differentiation^43–47^. We performed qRT-PCR analysis in whole colonic tissue to check for expression of CX3CR1 and CCR2. In fiber-high tissue as well as in muscle tissue, we found a significantly higher expression of CX3CR1 than CCR2 while in all other segments, expression levels of the two markers were comparable (Supplementary Fig. 2c). To check whether these changes resulted from altered MΦ populations, we investigated viable CD64^+^ MΦ by flow cytometry (Supplementary Fig. 3d). All MΦ expressed comparable levels of CD11c (Fig. 3f, upper graph). However, we found a significantly higher expression of CD14, an M2 marker^48^, in MΦ isolated from the mucosal lamina propria (LPMΦ) of fiber-high colonic tissue and in MΦ isolated from muscularis (MMΦ) (Fig. 3f, lower graph). MMΦ interact with neurons of the myenteric plexus and have been described as bipolarly shaped^49–51^. Using confocal fluorescence microscopy, we visualized a bipolar shape of LPMΦ in fiber-high segments similar to that seen with MMΦ (Fig. 3g, left and right image). In contrast, LPMΦ in fiber-low colonic segments exhibited a round or stellate phenotype (Fig. 3g, middle image). We found a significantly higher percentage of bipolar MΦ in fiber-high colonic tissue (Fig. 3g, bar graph). We mimicked the phenotype of the two colonic macrophage populations by differentiating blood-derived monocytes from healthy adult donors into M1 and M2 MΦ. After 10 days of differentiation, M2 MΦ showed an elongated bipolar shape (Supplementary Fig. 2d, e) and increased expression of CD14 and CX3CR1 (Supplementary Fig. 2f, g). In contrast, M1 MΦ exhibited a rounded shape and expressed moderate levels of CD14 while lacking CX3CR1 expression. To test the ability of the MΦ to differentiate T cells into certain effector T cells, we co-cultured M1 and M2 MΦ with autologous naïve CD4 T cells in the presence of anti-CD3 and performed a Th17 and Treg T cell differentiation assay. After 6 days of co-culture, the frequency of Th17 and Treg cells were analyzed and the ratio calculated. Th17/Treg ratio was significantly lower when bipolar CX3CR1^+^CD14^high^ M2 MΦ were used (Supplementary Fig. 2h). These observations suggested that fiber-high segments of HSCR colon harbor a bipolar CD64^+^CD11c^+^CD14^high^ MΦ population in the mucosa which correlates with decreased Th17 cell frequencies. Of note, due to the low viability of patient-derived colonic macrophages, no functional assays could be performed, and M2 MΦ did not fully resemble the colonic phenotype.

### Bipolar CD14^high^ MΦ colocalized with cholinergic fibers

Both MΦ and T cells express receptors for neurotransmitters, qualifying them for neuronal interaction^25,49,50,52–54^. Representative fluorescence images (20x and 63x) of fiber-high and fiber-low distal colonic tissue revealed CD64^+^ MΦ and CD3^+^ T cells in close neighborhood to AChE^+^ fibers (Fig. 4a). In the muscle layer, MΦ were only detected between thick cholinergic fibers. To identify the cell type interacting with mucosal nerve fibers, we performed fast-scanning confocal imaging using a Fiji software macro to determine the sum of overlapping pixel values (Fig. 3b and Supplementary Movie 1). Per patient (fiber-low segments, n=4; fiber-high segments, n=5; muscle, n=2), 100-200 cells were annotated, and the sum of pixel values was determined. The results showed that LPMΦ in fiber-high segments significantly colocalized with cholinergic fibers comparable to MMΦ, in contrast to LPMΦ of fiber-low segments (Fig. 4c). No significant overlap between CD3^+^ T cells and cholinergic fibers was found (Fig. 4d). Furthermore, neuron-associated LPMΦ in fiber-high tissue from HSCR patients mainly exhibited a bipolar shape (Fig. 4e). These results suggest that bipolar LPMΦ preferentially interacted with cholinergic fibers in aganglionic fiber-high tissue from HSCR patients.

**Fig. 4.**
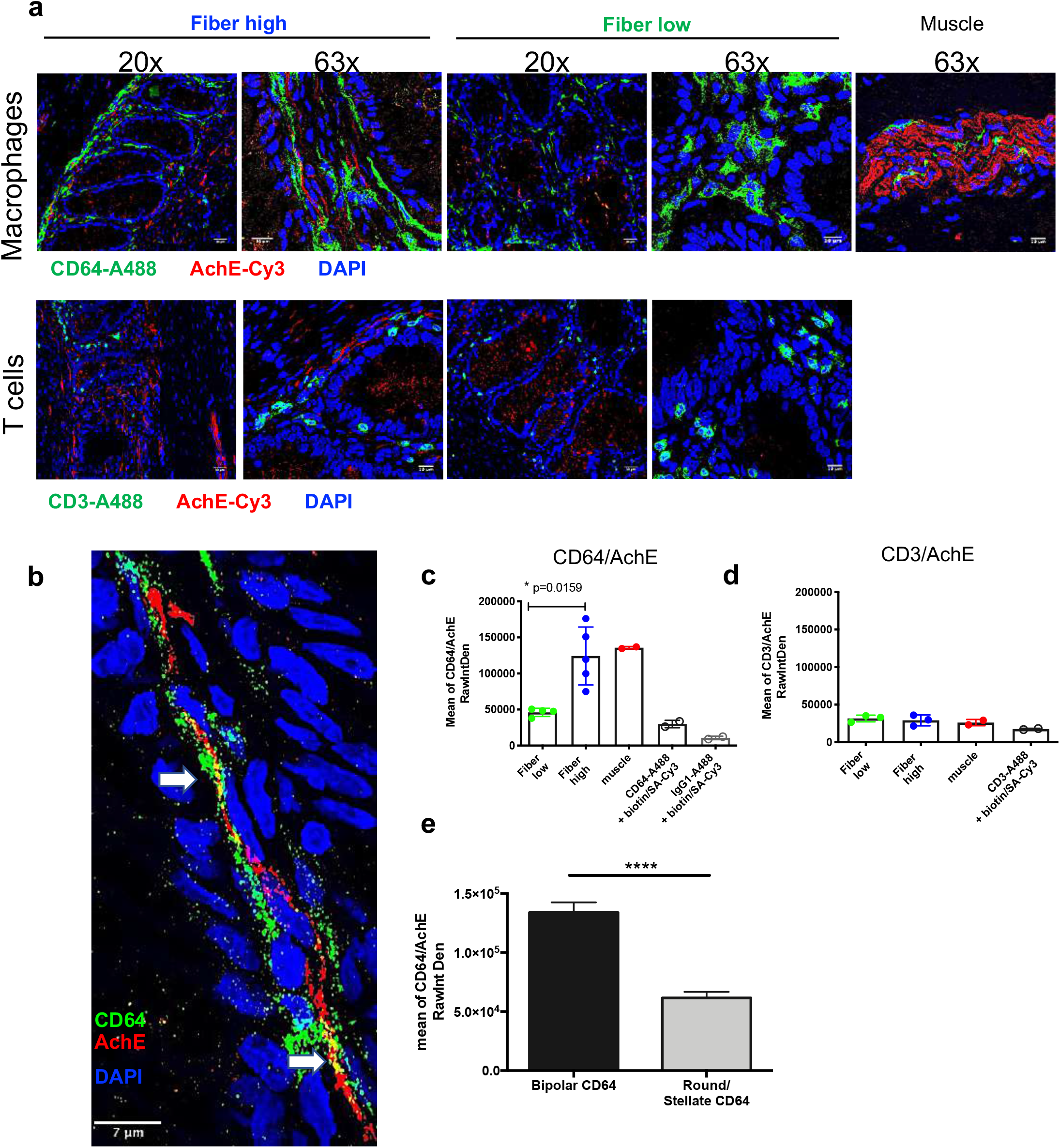
MΦ colocalize with cholinergic nerve fibers. **a**, Immunofluorescence of epithelial region in cryosections (5 μm, Swiss roll of rectum and sigmoid colon) of representative fiber-high and fiber-low patients. Anti-AChE (red); anti-CD64 or anti-CD3 (green); DAPI (blue). Image from muscle region (right) shows thick fiber bundles (AChE) together with CD64^+^ cells. Images were obtained by fluorescence (20x, scale bar 30 μm) and confocal (63x, scale bar 10 μm) microscopy and processed by Fiji software. **b**, Confocal immunofluorescence image of an intercrypt region from a fiber-high patient. Overlapping pixels (yellow, arrows) were visualized on a Z-projection image. See Supplementary movie. **c-d**, Quantification of the pixel intensity colocalization of CD64^+^ MΦ **(c)** or CD3^+^ T cells **(d)** with AChE^+^ fibers was performed by a Fiji software macro. Confocal images from distal colon regions of fiber-low (n=4 for CD64; n=3 for CD3) and fiber-high (n=5 for CD64; n=3 for CD3) patients as well as from muscle regions of HSCR patients (n=2) were taken. The following negative staining controls were used: CD64-A488 **(c)** or CD3-A488 **(d)** with biotin + streptavidin-Cy3 and IgG1-A488 + biotin/streptavidin-Cy3 (n=2). Per patient (dot), 200-300 cells were analyzed. Scatter plots show mean raw integrated density (RawIntDen) ± SEM. **e**, CD64^+^ MΦ from 3 fiber-high patients were investigated with respect to bipolar (n=170) or round/stellate (n=152) morphological phenotype. Raw integrated density of colocalized pixels between CD64 and AChE are shown. Significance was determined using unpaired nonparametric Mann-Whitney test (* p≤ 0.05; **** p≤ 0.0001). Source data are provided as a Source Data file.

### Cholinergic fibers suppressed Th17 inducing IL-23 expression in MΦ

We next sought to identify specific genes targeted by neuronal interaction. Therefore, we performed RNA sequencing in viable FACS sorted mucosal MΦ (CD45^+^HLA^+^CD64^+^) from fiber-low and fiber-high tissues. As control, we included MMΦ as well as blood-derived M1 and M2 MΦ differentiated *in vitro.* Principal component analysis (PCA) showed clear transcriptional separation of blood-derived M1 and M2 MΦ, whereas colonic LPMΦ and MMΦ accumulated together (Fig. 5a). Macrophage-specific genes (CD64, CD14, ITGAM, CX3CR1, CCR2, ITGAX, and CSF1R) were similarly expressed between colonic populations while MMΦ expressed muscle-specific Lyve1 (Fig. 5b)^55^. When testing for gut-specific M2 markers (AhR, CD200R1, AREG, CD209, TGF-β2, and IL-10), we found elevated expression in colonic subsets similar to blood-derived M2 MΦ. In contrast to LPMΦ from fiber-low segments, LPMΦ from fiber-high tissue as well as MMΦ showed decreased expression of chemokine receptor CCR7 crucial for lymph-node homing, indicating a tissue-resident population (Fig. 5b). Differential gene expression analysis yielded 24 significantly changed genes between LPMΦ isolated from fiber-low and fiber-high colonic tissues (Fig. 5c, d). Strikingly, IL-23a, a subunit of the Th17-inducing cytokine IL-23, was significantly downregulated in LPMΦ from fiber-high segments, suggesting a correlation to cholinergic innervation. Additionally, we found a significant reduction of IL-6 and CCL-20, both of which are involved in Th17 differentiation and recruitment, but no significant change in TNF-α (Fig. 5e).

**Fig. 5.**
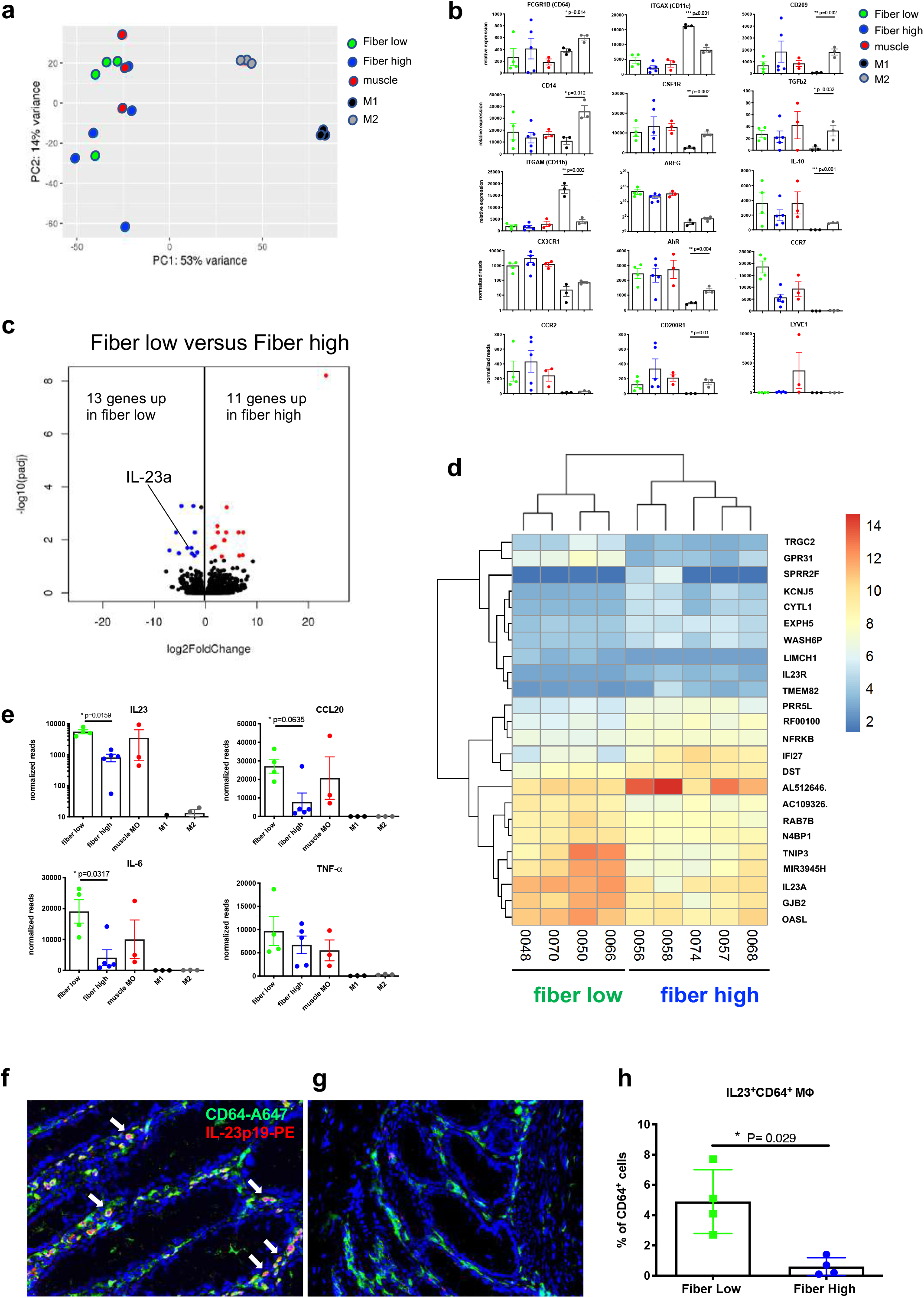
Fiber-associated MΦ express diminished IL-23 levels. RNA sequencing was performed on sorted viable colonic MΦ (CD45^+^HLA^+^CD64^+^, Supplementary Fig.3d and Methods) and blood-derived M1 and M2 MΦ. **a**, PCA plot shows similarity between the different populations based on their first two principal components. The top 500 genes, selected by highest row variance, were used. PC, principal component. Fiber-low (green, n=4); fiber-high (blue, n=5); muscle (red, n=3); M1 (black, n=3); M2 (grey, n=3). **b**, Normalized sequence reads of selected macrophage-related genes. Scatter plots show means ± SEM. Significance was determined using unpaired nonparametric Mann-Whitney test or unpaired t-test (for M1/M2 MΦ) (* p≤ 0.05; ** p≤ 0.01; *** p≤ 0.001) **c**, Global transcriptional changes between macrophages isolated from fiber-low colonic tissue and fiber-high colonic tissue visualized by a volcano plot. Genes with an adjusted p-value < 0.05 and log2 fold change greater than 1 (red dots) and less than −1 (blue dots) are indicated. **d**, Heat map of differentially expressed genes (from b) between MΦ in fiber-low colonic tissue and fiber-high colonic tissue. Each column represents one patient. **e**, Normalized sequence reads of selected genes. Scatter plots show means ± SEM. Significance was determined using unpaired nonparametric Mann-Whitney test or unpaired t-test (for M1/M2 MΦ) (* p≤ 0.05; ** p≤ 0.01; *** p≤ 0.001). **f-g**, IL-23^+^CD64^+^ MΦ were assessed by immunofluorescence in 5 μm colonic cryosections. A representative image of mucosal/epithelial region from fiber-low tissue (**f**) and fiber-high tissue (**g)** is shown. Scale bar 30 μm. CD64 (A647, green); IL23p19-PE (red) **h**, Quantitative analysis of IL-23 expression in macrophages evaluated in images from 4 fiber-high and 4 fiber-low patients. A total of 900-4700 CD64^+^ cells per patient were analyzed for IL-23 expression and the percentage of IL23^+^CD64^+^ cells (% of total macrophages) calculated. Scatter plots show means ± SEM. Significance was determined using unpaired nonparametric Mann-Whitney test (* p≤ 0.05). Source data are provided as a Source Data file.

Using immunofluorescence analysis, we quantitatively analyzed IL-23 protein expression in CD64^+^ LPMΦ in cryosections from fiber-low and fiber-high colonic tissue (Fig. 5f, g). LPMΦ in 4 to 8 regions per patient were analyzed and quantified using CellProfiler software. Fiber-low colonic tissue showed significantly increased frequencies of IL-23-positive LPMΦ compared to fiber-high colonic tissue (Fig. 5h). This result indicated that the lack of mucosal cholinergic fibers correlated with increased expression of the Th17 inducing cytokine IL-23 in LPMΦ.

### Distinct expression of macrophage-neuron crosstalk transcripts

Next, we aimed to identify receptors possibly involved in the neuronal modulation of LPMΦ. Based on transcriptional analysis, we verified the expression of receptors for neurotransmitters, neuropeptides, neurotrophic factors, and axon-guidance proteins (Fig. 6). Considering the crucial role of α7nAChR in mediating the CAIP, we found elevated levels of the receptor in LPMΦ isolated from fiber-high segments from HSCR patients (Fig. 6a and f). In contrast, other receptors reported to be involved in cholinergic immune suppression such as receptors for serotonin^56^, catecholamine (ADRB2)^50,57^, or neuropeptides (NMUR1, VIPR1, SSTR2, and CALCRL)^58–65^ were not upregulated in fiber-high LPMΦ compared to fiber-low LPMΦ or were not expressed (NMUR1) (Fig. 6b, c and f). MΦ secrete different neurotrophic and axon-guidance proteins as well as respective receptors. Fiber-associated LPMΦ expressed growth/differentiation factor (GDF) 15 and, to a lesser extent, GDF11 (Fig. 6d and f). Within the family of axon-guidance proteins, we found elevated expression of the receptors neuropilin-1 (NRP1), UNC-5 homology B (UNC5B), and ephrin type B-receptor 2 (EPHB2) binding semaphorin 3A (SEMA3A), netrin, and ephrin-B family members, respectively (Fig. 6e and f). In contrast, LPMΦ isolated from fiber-low segments showed elevated expression of SEMA3A and SEMA7A compared to LPMΦ from fiber-high segments (Fig. 6e and f).

**Fig. 6.**
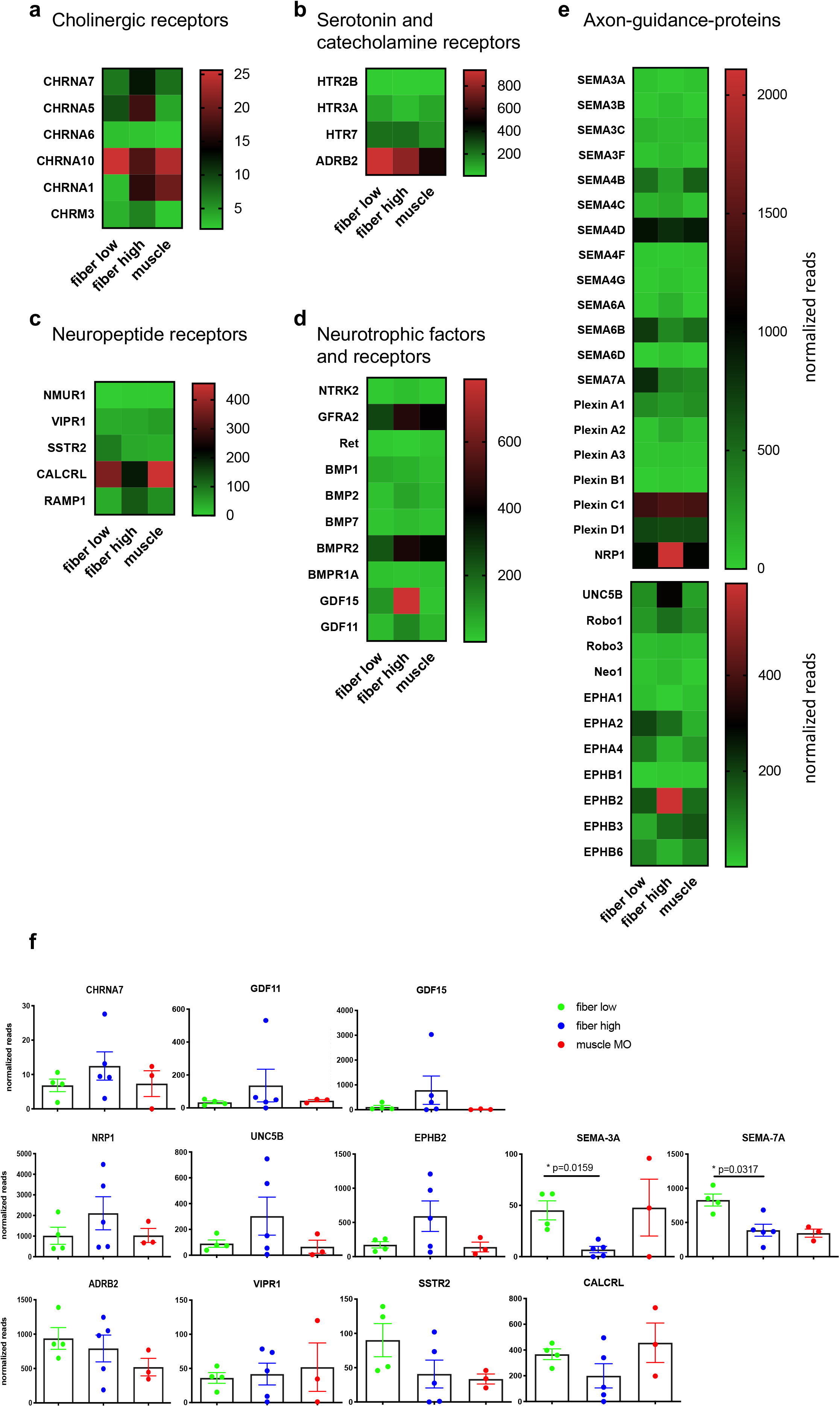
Receptors involved in macrophage-neuron crosstalk. **a-e**, Heat maps of selected genes possibly involved in macrophage-neuron crosstalk. Normalized reads are shown. RNA sequencing was performed on sorted viable colonic MΦ (CD45^+^HLA^+^CD64^+^) from mucosa of fiber-low (n=4) and fiber-high (n=5) colonic tissue as well as muscle tissue (n=3). Heat maps show mean values of each group. **a**, Expression of nicotinic (CHRNA) and muscarinic (CHRM) ACh receptors. **b**, Expression of serotonergic (HTR) and catecholaminergic (ADRB) receptors. **c**, Expression of receptors for neuromedin U (NMUR1), vasoactive-intestinal-peptide (VIPR1), somatostatin (SSTR2), calcitonin-gene-related peptide (CALCRL/RAMP1). **d**, Expression of neurotrophins (bone-morphogenic proteins [BMP] and growth/differentiation factors [GDF]) and neurotrophin receptors (NTRK2 binds brain-derived-neurotrophic factor (BDNF) and neurotrophin (NT)-4; GFRA2 and RET bind GDNF and neurturin. **e**, Expression of axon-guidance proteins and respective receptors. Semaphorins (SEMA) bind to plexins or neuropilin-1 (NRP1). UNC-5 represent the receptor for netrin. Receptor proteins ROBO1 and 3 bind slit proteins. Neogenin (Neo) receptor binds to repulsive guidance protein (RGM). Ephrins bind to Eph receptors EPHA and EPHB. **f**, Normalized sequence reads of selected genes. Scatter plots show means ± SEM. Significance was determined using unpaired nonparametric Mann-Whitney test (*p≤ 0.05). Source data are provided as a Source Data file.

Overall, these data suggest a possible involvement of α7nAChR in neuron-MΦ crosstalk. In addition, fiber-associated MΦ seemed to support neuronal growth and differentiation via GDF production and axon-guidance proteins. However, data analysis was limited by low sample numbers and high variance and thus needs further validation.

### Microbial dysbiosis in colonic fiber-low tissues from HSCR patients

Lacking intestinal MΦ or ENS affects the composition of intestinal microbiome^66–69^. In a chemically induced colitis model in mice, CX3CR1^+^ MΦ were crucial to maintain the intestinal barrier function to prevent bacterial translocation^47,70^. To evaluate if the presence of cholinergic fibers impacts on bacterial translocation, we performed 16S rDNA FISH analysis of fiber-low and fiber-high cryosections. Bacteria translocated to lamina propria were detected in both groups, but no significant difference was found (Supplementary Fig. 4a, b). We then checked the composition of microbial communities by 16s rDNA sequencing in cryosections of fiber-low, fiber-high, and ganglionic CD tissue from HSCR patients. Alpha diversity analysis showed an increase in operational taxonomic units (OTUs) in fiber-low tissue when compared to ganglionic CD tissue, whereas estimated OTU richness as well as beta diversity were comparable in all groups (Fig. 7a and Supplementary Fig. 4c). On the phylum levels, *Actinobacteria* seemed to be less prevalent in aganglionic tissue compared to ganglionic tissue (Fig. 7b). Within the bacterial family compositions, *Actinobacteria* were significantly less prevalent in fiber-low segments compared to ganglionic CD, whereas other families were similarly expressed (Fig. 7c). The main representative species of the *Actinobacteria* were several *Bifidobacteria (B. longum* (OTU1)*; B. bifidum* (OTU6)*; B. pseudocatenulatum* (OTU5), with *B. longum* showing the highest frequency in all samples (Supplementary Table 5). Differential bacterial family analysis between fiber-low and fiber-high colonic tissue revealed a significant increase in *Staphylococcaceae* in fiber-low colonic segments, whereas in fiber-high tissue, *Enterobacteriaceae* and *Clostridiales Family_XI* were upregulated (Fig. 7c). Within the *Staphylococcaceae* family, *S. aureus* (OTU7) was the only detectable species within the core microbiome (Supplementary Table 5). The *Clostridiales Family_XI* was represented by 4 genera, i.e., *Finegoldia* (OTU825)*, Peptoniphilus* (OTU196)*, Anaerococcus* (OTU543) and *Ezakiella* (OTU27, OTU63). The main representative species of *Enterobacteriaceae* family was *Escherichia coli* (OTU2), which represented the second frequent bacterial species from the core microbiome (Supplementary Table 5). These observations suggested a different composition of colonic translocated bacteria between fiber-low and fiber-high tissue including a decrease in *Actinobacteria* and prevalence of *Staphylococcaceae* family in fiber-low segments lacking cholinergic mucosal innervation.

**Fig. 7.**
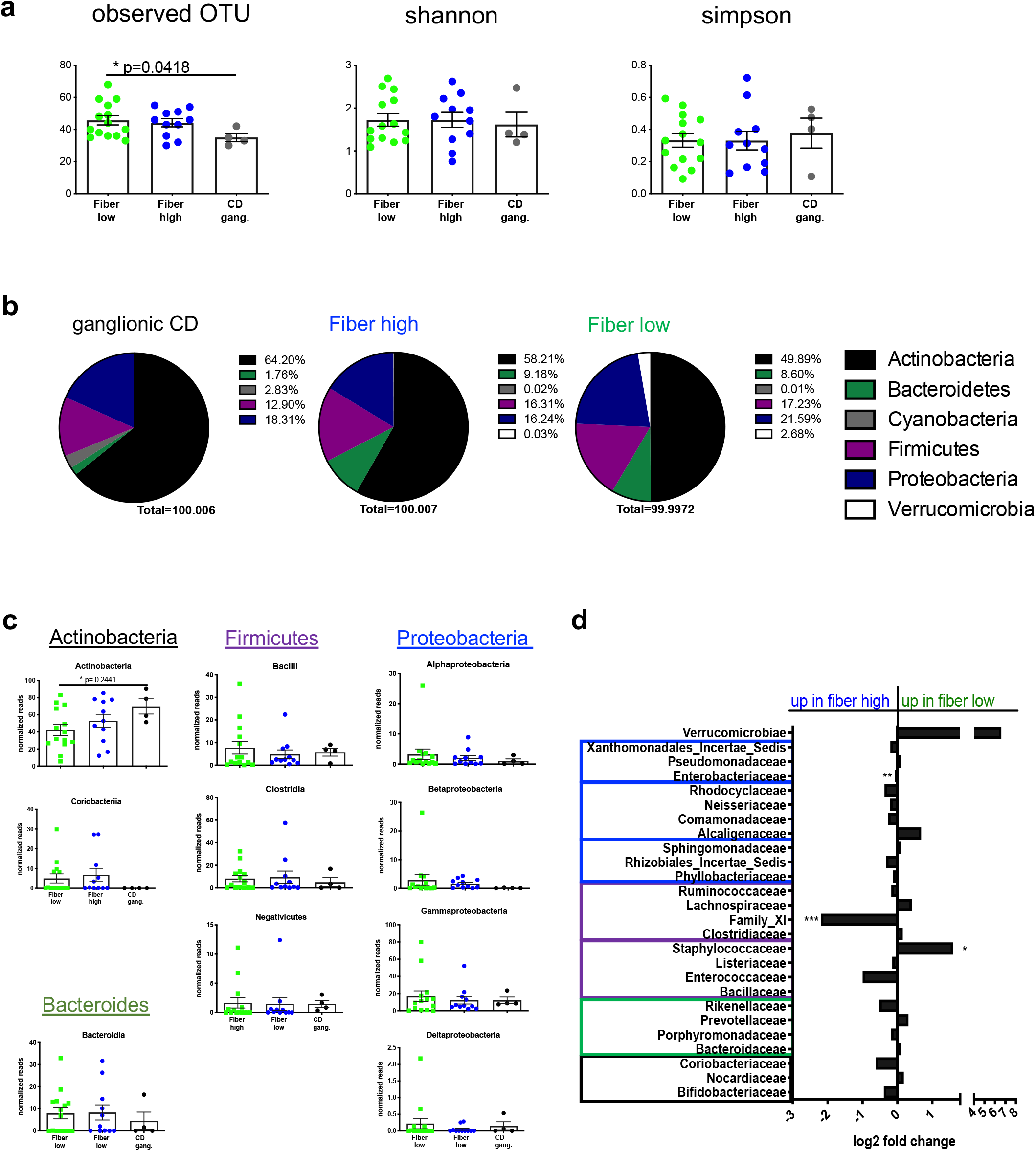
Colonic microbial dysbiosis in fiber-low colonic tissue from HSCR patients. **a-d,** 16S rRNA analysis of colonic tissue from aganglionic (fiber-low n=14; fiber-high n=11) and ganglionic (CD gang. n=4) segments of age-matched HSCR patients. **a**, Alpha diversity analysis show effects on observed operational taxonomic units (OTUs) and estimated OTU richness by Shannon and Simpson indices. **b**, Pie charts show microbial composition in each group on phylum level. **c**, Scatter plots show normalized reads of bacterial classes dedicated to respective phyla. Scatter blots show mean values ± SEM. Significance was determined using unpaired nonparametric Mann-Whitney test (* p≤ 0.05). **d**, Differential bacterial family analysis of fiber-low versus fiber-high groups. Asterisk mark significantly changed families with an adjusted p value < 0.05 (***p= 4.01×10^-15^; **p= 2.08×10^-4^; *p= 0.022). Families are assigned to respective phyla by color code. Source data are provided as a Source Data file.

### HSCR patients with fiber-low phenotype showed a higher incidence of posto-perative enterocolitis

Next, we evaluated the impact of fiber phenotype on the development of postoperative enterocolitis. One year after pull-through surgery, 42 HSCR patients were followed up, and postoperative enterocolitis manifestation was evaluated from medical records. Enterocolitis was defined to fulfill at least 3 of the following criteria: diarrhea (>3x/day), vomiting, temperature >38.5°C, antibiotics, hospitalization, meteorism, and high inflammatory blood parameters (i.e., leucocytes, CRP). Anthropometric, clinical, environmental, maternal, and dietary factors, collected from the parent questionnaire, were analyzed for potential enterocolitis risk factors. A total of 9 (of 42) patients developed enterocolitis one year after surgery (Table 1a). Development of enterocolitis was significantly associated with cesarean delivery at birth of the infant (p=0.012). Seven of the 9 patients exhibited a fiber-low and only two a fiber-high phenotype, showing a higher incidence of fiber-low phenotype among the children who developed enterocolitis (78% versus 22%, p=0.634). To validate the observations, we analyzed a retrospective HSCR patient cohort collected between 2003 and 2018 at the UKBB (Supplementary Table 6). A total of 29 HSCR patients were included. We did not observe any significant patient bias in terms of anthropometric or clinical factors between fiber-low and fiber-high tissues (Supplementary Table 7). Enterocolitis development was recorded in 14 of 29 patients but showed no correlation with anthropometric or clinical data (Table 1b). Strikingly, enterocolitis development correlated significantly with the fiber phenotype showing a higher incidence in children with the fiber-low phenotype (86%, 12/14 children) compared to the fiber-high phenotype (14%, 2/14). These observations suggest that HSCR patients with fiber-low phenotype might have a higher risk to develop enterocolitis and that the fiber phenotype could serve as a prognostic marker for risk of enterocolitis development.

**Tab. 1.** Risk factors for post-operative enterocolitis. **a,** Prospective HSCR patient cohort (n=42). Values are median (range) or number (percentage); (n): number of data available; length of aganglionosis: rectum (1), sigmoid colon (2), colon descendens (3), colon transversum (4), colon ascendens (5). Differences between enterocolitis development (yes/no) were tested with the Wilcoxon rank-sum test and χ2-test, as appropriate. No follow-up for patients no. 0023 and 0032 available. **b**, Retrospective HSCR patient cohort (n=29). Values are mean (SD), median (range), or number (percentage); n: number; SD: standard deviation. Differences between enterocolitis development (yes/no) were tested with the Wilcoxon rank-sum test and χ2-test, as appropriate. Source data are provided as a Source Data file.

| Prospective HSCR patient cohort | Enterocolitis |  | p-value |
| --- | --- | --- | --- |
| Anthropometric data and potential risk factors, (n) | Yes n=9 | No n=33 |  |
| Age at operation, months (42) | 3 (1–127) | 5 (2–70) | 0.211 |
| Sex, male (42) | 7 (78) | 27 (81) | 0.784 |
| Preterm birth (38) | 2 (22) | 1 (3) | 0.068 |
| Operation procedure trans-anal (42) | 8 (89) | 29 (88) | 0.934 |
| Length of aganglionosis (42) | 1 (1-5) | 2 (1-5) | 0.061 |
| Pets (37) | 1 (11) | 11 (39) | 0.116 |
| Patient living in rural area (40) | 3 (33) | 17 (55) | 0.256 |
| Antibiotic treatment (38) | 2 (22) | 3 (10) | 0.357 |
| Siblings (39) | 8 (89) | 22 (73) | 0.331 |
| Breastfed at operation (39) | 9 (100) | 27 (90) | 0.323 |
| Maternal age at delivery, years (35) | 27 (23-39) | 29 (21-43) | 0.788 |
| Parental smoking (38) | 4 (44) | 5 (17) | 0.094 |
| Caesarean Section (38) | 6 (75) | 8 (27) | 0.012 |
| Maternal Allergy (38) | 1 (11) | 8 (31) | 0.236 |
| Fiber low (30) | 7 (78) | 23 (70) | 0.634 |
| Fiber high(12) | 2 (22) | 10 (30) | 0.634 |

b
| Retrospective HSCR patient cohort | Enterocolitis |  | p-value |
| --- | --- | --- | --- |
|  | Yes (n=14) | No (n=15) |  |
| Age at operation, months | 4 (1–48) | 9 (1–171) | 0.229 |
| Sex, male | 11 (79) | 14 (93) | 0.249 |
| Preterm birth | 1 (7) | 1 (7) | 0.960 |
| Antibiotic treatment | 10 (7) | 11 (73) | 0.909 |
| Caesarean Section | 7 (50) | 5 (33) | 0.124 |
| Fiber low | 12 (86) | 3 (20) | <0.001 |
| Fiber high | 2 (14) | 12 (80) | <0.001 |

## Discussion

In recent years, multiple studies mainly using different mouse disease models demonstrated the reciprocal relationship between immune and nervous systems. In humans however, difficulty to study individual effects of neurotransmitters leads to a lack of studies providing direct evidence of neuronal-immune crosstalk. Here, we provided unique evidence of a cholinergic neuron-macrophage crosstalk in the colon correlated with reduced type 17-mediated immune reactions. By studying colonic tissue of HSCR patients characterized by the lack of an ENS and extramural mucosal cholinergic innervation, we deciphered the relationship between neuronal ACh and the mucosal immune system. Using confocal colocalization studies, we identified elongated tissue-resident MΦ interacting with cholinergic nerve fibers. Differently shaped intestinal MΦ populations were described before. Gabanyi et al. described bipolar-shaped MMΦ with a tissue-protective anti-inflammatory M2 phenotype contrary to stellate-shaped LPMΦ^50^ The elongated profile allows MMΦ to interact and wrap around neighboring neurons. At steady state, MMΦ regulate ENS functions and therefore gut motility by providing bone morphogenetic protein (BMP) 2 in response to microbial stimuli^49^. Tamoxifen-induced deletion of CX3CR1^+^ tissue-resident MΦ leads to loss of enteric neurons^71^. Interestingly, MMΦ develop independently of the ENS^71,72^. The ENS supplies colony-stimulating factor1 (CSF1) needed for M2 MΦ differentiation and survival^49,73^. Upon *Salmonella typhimurium* infection, MMΦ interact with sympathetic adrenergic neurons via β2 adrenergic receptors to induce a tissue- and neuron-protective gene expression program^50^. In a model of POI which is characterized by surgery-induced inflammatory response of MMΦ and impaired motility, VN stimulation or pharmacologic triggering of cholinergic enteric neurons prevented MΦ activation and reduced POI in a α7nAChR-dependent manner in both mice and humans^56^. However, anti-inflammatory neuro-immune modulation seems to involve different neuronal receptors in type 2-mediated intestinal disease models. For example, in a murine model of food allergy, VN stimulation ameliorated disease severity independently of α7nAChR^74^. Uptake of food antigens by anti-inflammatory CX3CR1^+^ MΦ and subsequent induction of foodspecific Tregs effectively prevent allergic reactions^44^. VN stimulation increases phagocytic activity in CX3CR1^+^ MΦ, independently of α7nAChR, but phagocytic activity remains dependent on α4β2nAChR^75^. Using a murine helminth infection model, the Artis group demonstrated that the cholinergic neuropeptide neuromedin U activates ILC2 thus increasing the protective immune response and parasitic clearance^58,59^. In contrast, signals from sympathetic adrenergic neurons lead to attenuated ILC2 effector function and prevent chronic pathologic type 2 inflammation via activation of ILC2-specific β2 adrenergic receptors^76^. In our study, IL-23, a cytokine promoting the maintenance of Th17 cells^77^, is significantly increased in LPMΦ isolated from fiber-low tissue lacking cholinergic innervation compared to innervated fiber-high tissue. Consistent with this finding, elevated Th17 cells were detected in fiber-low colonic tissue compared to fiber-high colonic tissue. Th17 cells are indispensable for the clearance of extracellular bacteria and fungi and therefore for protective immunity. Dysregulation of cytokines controlling Th17 cells predisposes to the development of autoimmune diseases and inflammatory bowel disease^36,78,79^. Genome-wide studies identified genes, including the gene for IL-23 receptor, involved in Th17 differentiation associated with a susceptibility to inflammatory bowel disease^78,80–82^. IL-23 is produced by dendritic cells and MΦ in response to extracellular bacteria and fungi recognized by special innate patternrecognition receptors, the toll-like receptor (TLR) 2 and nucleotide oligomerization domain containing protein 2 (NOD2)^83,84^. Under inflammatory conditions, monocyte-derived CCR2^+^CX3CR1^+^ MΦ infiltrate the gut mucosa and prime Th17 cell responses^43,47^. At steady state, CX3CR1^+^ MΦ are indispensable for epithelial integrity, control of bacterial translocation, and development of oral and commensal tolerance by secreting anti-inflammatory cytokines and maintaining Treg^44,68^. CX3CR1 MΦ further promote intestinal homeostasis by microbiota-dependent crosstalk with IL-22-secreting ILC3^85^.

In our study, we found higher tissue expression of CX3CR1 than CCR2 in fiber-high colonic segments of HSCR patients and identified two MΦ populations based on CD14 expression and shape. CD14^+^ bipolar MΦ were found in close proximity to nerve fibers in both muscularis and lamina propria, expressing low levels of CCR7, suggesting a tissue-resident phenotype^86^. CD14^low^ stellate-shaped MΦ were present in non-innervated lamina propria tissue, expressing CCR7 indicating a migratory population. Based on their transcriptional profile, LPMΦ isolated from both fiber-low and fiber-high colonic tissues exhibit a gut-imprinted M2 phenotype (CX3CR1, AREG, AhR, CSF1R, CD200R, CD209, TGFβ2, and IL-10), but they differed in Th17 cytokines (IL-23 and IL-6) and Th17 chemokines (CCL20). Consequently, cholinergic signals seemed to be associated with reduced type 17 immune responses. In a murine model of psoriasis, sensory nociceptive neurons augment dermal dendritic cell-mediated IL-23 production^64^. The neurons directly sense disease-inducing *Candida albicans* and release the neuropeptide calcitonin gene-related peptide (CGRP) which increases IL-23 production^65^. Surprisingly, another study uncovered an immunosuppressive effect of nociceptive fibers during *S. aureus* infection indicating a role for the properties of the underlying pathogen or target immune cell on nerve fiber-induced immune modulation^87^.

Neurotrophic factors and their respective receptors are crucial for ENS development and function. In fact, the underlying genetic defects in HSCR patients involve signaling pathways controlling migration and development of the ENS, including the two major HSCR genes, rearranged during transfection (Ret) and endothelin receptor type B (EndrB)^88,89^. In a study using Ret-deficient mice, the authors demonstrated that microbiota-induced glial cell-derived neurotrophic factors induce IL-22 in Ret-expressing ILC3 therefore maintaining intestinal homeostasis^90^. Ret expression was not detected in colonic MΦ populations, and we did not detect any changes in ILC3 frequencies between fiber-low and fiber-high colonic segments. Here, we used RNA-seq transcriptional analysis to identify factors and receptors involved in neuro-immune interactions. However, the RNA-seq data were highly variable within the groups and were not validated in qPCR analyses or on protein levels. Further studies are needed to fully uncover receptors and mechanisms involved in neuron-MΦ interactions. Besides the contribution of cholinergic receptors, we supposed the contribution of neurotrophic factors, axon-guidance proteins, and their respective receptors in neuron-MΦ interactions. GDF15 has been reported to suppress inflammatory responses in a murine model of endotoxin-induced sepsis and prevent maturation of inflammatory dendritic cells in a heart-transplantation model^91,92^. Using an imiquimod-or IL-23-induced mouse model of psoriasis-like skin inflammation, GDF11 administration attenuated the severity of skin inflammation via suppression of the NF-κB signaling pathway^93^. Recently, NRP-1 was identified in adipose-tissue MΦ protecting from metabolic syndrome by favoring fatty acid uptake and β-oxidation away from pro-inflammatory M1 polarization and glycolysis^94^. Additionally, in a model of a neurodevelopmental disorder, brown adipose-tissue MΦ control sympathetic tissue innervation critical for thermogenesis via the plexin-semaphorin pathway^52^.

Inflammation in chemically induced colitis is suppressed by administering recombinant netrin-1^95^ whereas UNC5B deficiency exacerbates chemically induced colitis^96^. SEMA3A regulates neurite growth cone repulsion suggesting a role of LPMΦ from fiber-low tissues in the inhibition of mucosal innervation^97,98^. Besides its crucial role in neurogenesis, SEMA7A supports the generation of monocyte-derived inflammatory cytokines and differentiation of inflammatory T cells^99,100^. More studies are needed to evaluate the involvement of the different receptors in our study.

The microbial composition of feces from HSCR patients with enterocolitis is shifted towards opportunistic pathogens such as *Candida albicans* und *Clostridium difficile^1,10,101,102^.* In our study, none of the patients was diagnosed with enterocolitis at the time of the pull-through surgery. However, higher baseline frequencies of Th17 cells and IL-23 might promote *C. difficile* infection, as was recently postulated^103^. Microbial prevalence and composition were altered in aganglionic tissue from fiber-low and fiber-high phenotypes compared to ganglionic segments. *Actinobacteria* were significantly reduced in fiber-low segments compared to ganglionic CD tissue. Compared to fiber-high colonic tissue, fiber-low tissue is associated with higher levels of *Staphylococcus aureus* previously reported to be associated with an increased Th17 response^104^. In contrast, *Clostridiales Family_XI* was upregulated more than 2 log2 fold in fiber-high tissue compared to fiber-low tissue, which was associated with a decreased risk of *Clostridium difficile* infection and necrotizing enterocolitis in preterm children^105,106^. In our study, differential bacterial analysis was limited due to the small patient samples per group and high variability within the groups. The fact that we analyzed bacterial communities in colonic tissue and rather than feces further limited the data analysis.

Overall, 9 of 42 patients developed HSCR-associated enterocolitis one year after pull-through-surgery. Interestingly, development of enterocolitis was significantly associated with cesarean delivery at birth of the infant in the prospective HSCR patient cohort. Microbial composition in children delivered by cesarean section is known to be associated with skin commensals such as *Staphylococcus,* while the microbiome in children delivered by natural birth mode is characterized by vaginal bacteria, *e.g., Bifidobacteria* and *Lactobacillus^107–110^.* However, since cesarean section was not a significant risk factor in the retrospective patient cohort, its role in enterocolitis development needs to be further examined. Analyzing the correlation of fiber phenotype with enterocolitis development one year after pull-through surgery showed a higher incidence for fiber-low (78%) phenotype than fiber-high (22%) phenotype in patients developing enterocolitis. The result was validated in a retrospectively analyzed HSCR patient cohort and showed a significant incidence of 86% (fiber-low phenotype) versus 14% (fiber-high phenotype) to develop enterocolitis. In the retrospective cohort, fiber-low and fiber-high phenotypes were equally distributed (15 versus 14). Almost 50% of the study population developed enterocolitis compared to 21% in the prospective cohort.

HSCR represents a rare disease, and low patient numbers limit data analysis. However, we anticipate validating our results in additional retrospective HSCR patient cohorts. Interestingly, in HSCR mouse models using EndrB-deficient mice, no mucosal nerve fibers were detected, and the mice exhibited microbial dysbiosis leading to enterocolitis^111–113^. The absence of cholinergic fibers could therefore serve as a predictive marker for enterocolitis and improve personalized patient management. With the current post-operative therapeutic management, clinicians can react to inflammatory bowel situations only at the time point of enterocolitis onset. Using the fiber phenotype as a prognostic marker for the risk of enterocolitis, clinicians could directly react after pull-through surgery to prevent postoperative enterocolitis. Possible therapies include high-volume enemas, antibiotic interval treatment, pre- and probiotic therapy, as well as dietary adaptations. Additionally, interventional surgical corrections may be considered. For instance, it is still unclear how the neuronal phenotype of a resected colon segment might predispose for enterocolitis development. It has been reported that colonic inflammatory Th17 cells migrate to neighboring mesenteric lymph nodes, triggering enterocolitis development. Because the mesentery is generally not removed during surgery, this could explain why patients may develop enterocolitis after removal of affected aganglionic colon segments. Using the fiber phenotype as a prognostic enterocolitis marker would enable possible resection of the neighboring mesentery lymph nodes/tissue as a strategy to prevent enterocolitis. Further, immunotherapies targeting IL-17 or IL-23 rather than TNF-α could open new therapeutic treatment options.

Using a HSCR patient cohort lacking the ENS, we provided evidence that extrinsic mucosal cholinergic neurons interacted with MΦ and correlated with reduced type-17 immune response most likely mediated by reduced MΦ-dependent IL-23 expression. HSCR patients lacking mucosal extrinsic cholinergic neurons had a higher risk of developing postoperative enterocolitis. The mucosal fiber phenotype could serve as a new prognostic marker for HSCR-associated enterocolitis risk and may offer a personalized patient care and future therapeutic options.

## Author contributions

S.K.: study design; design, execution, and supervision of the experiments; data acquisition, analysis, and interpretation; writing and revision of the manuscript; V.G-F.: execution of the experiments; data acquisition and analysis; contribution to manuscript drafting; U.K.: execution of the experiments; data acquisition and analysis; M.M.: execution of the experiments; data acquisition and analysis of retrospective patient cohort; J.U.: statistical analysis; E.B.: acquisition of patient samples; revision of the manuscript; U.S., S.J.T., T.K., E.S, P.S., D.C., P.R., B.R., and NIG Study Group: recruitment, characterization, and follow-up of patients; revision of the manuscript S.H-C.: study design, recruitment, characterization, and follow-up of patients; revision of the manuscript.

## Abbrevations used in this paper

HSCR: Hirschsprung’s disease
FACS: Fluorescence-activated cell sorting
qRT-PCR: Quantitative reverse-transcribed real-time polymerase chain reaction
IL: Interleukin
Th17: T-helper-17
ENS: Enteric nervous system
ACh: Acetylcholine
α7nAChR: α7-nicotinic ACh receptor
CAIP: Cholinergic-anti-inflammatory-pathway
VN: Vagus nerve
MΦ: Macrophages
MMΦ: Muscle macrophages
TNF: tumor necrosis factor
POI: Postoperative ileus
CD: Colon descendens
REC: Rectum
SC: Sigmoid colon
VIP: Vasoactive intestinal peptide
TH: Tyrosine hydroxylase
NOS: Nitric oxide synthase
NK: Natural killer
NCR: NK cell receptors
APC: Antigen-presenting cell
PCA: Principal component analysis
HLA: Human leucocyte antigen
GDF: Growth/differentiation factor
NRP1: Neuropilin-1
UNC5B: UNC-5 homology B
EPHB2: Ephrin type B-receptor 2
SEMA: Semaphorin
LP: Lamina propria
FISH: Fluorescence in situ hybridization
OTUs: Operational taxonomic units
BMP: Bone morphogenetic protein
TLR: Toll-like receptor

## Acknowledgment

We thank all the patients and their parents as well as volunteer blood donors for their contribution. We also thank T. Lopes, E. Traunecker, and L. Raeli at the Flow Cytometry Facility of the Department of Biomedicine (Basel) for cell sorting, A. Loynton-Ferrand, K. Schleicher, and O. Biehlmaier at the Imaging Core Facility of the Biocenter (Basel) for assistance in confocal microscopy, D. Calabrese at the Histology Facility of the Department of Biomedicine (Basel) for histological support, P. Weigold at Microsynth AG (Balgach) for microbiome analysis, and S. Rogers at MediWrite (Basel) for her excellent manuscript revision.

---

**Supplementary Table 1:**
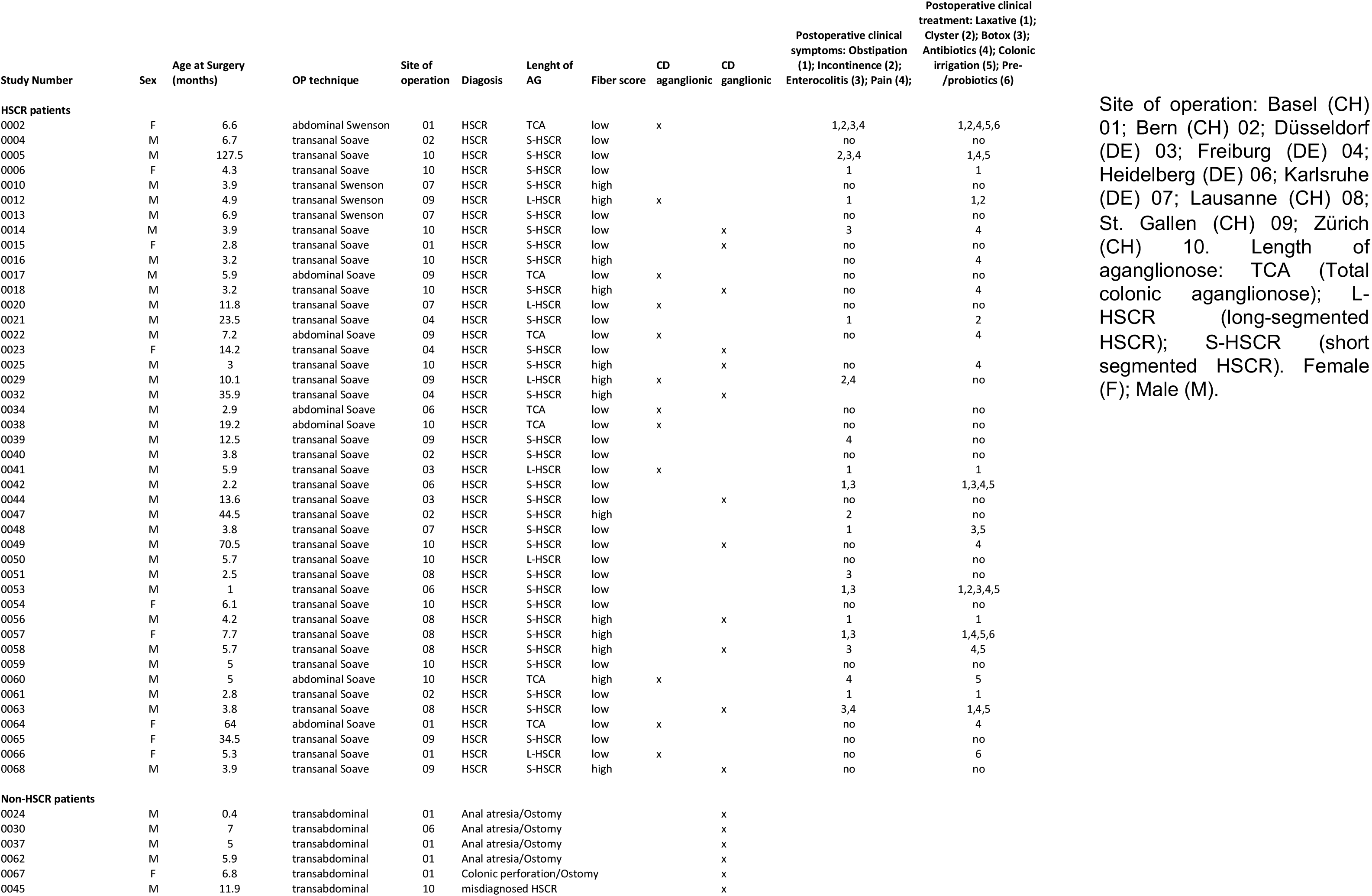
Overview of prospective HSCR patient cohort

**Supplementary Table 2:** Enzyme histochemistry performed at the different pathology units of recruiting clinics used for HSCR diagnosis

| Pathology unit | Detection of ganglia | Detection of extrinsic nerve fibers |
| --- | --- | --- |
| Basel, Freiburg, Karlsruhe | LDH, NOS, SDH | AChE |
| Zürich | LDH | AChE |
| Bern | LDH, NOS | AChE |
| St. Gallen | SDH | AChE |
| Lausanne | LDH | AChE |
| Heidelberg | Calretinin | AChE |
| Düsseldorf | NADPH, LDH | AChE |

Lactic dehydrogenase (LDH), succinic dehydrogenase (SDH), nitroxide synthase (NOS) Nicotinamide adenine dinucleotide (NADPH), acetylcholinesterase (AChE)

**Supplementary Table 3:**
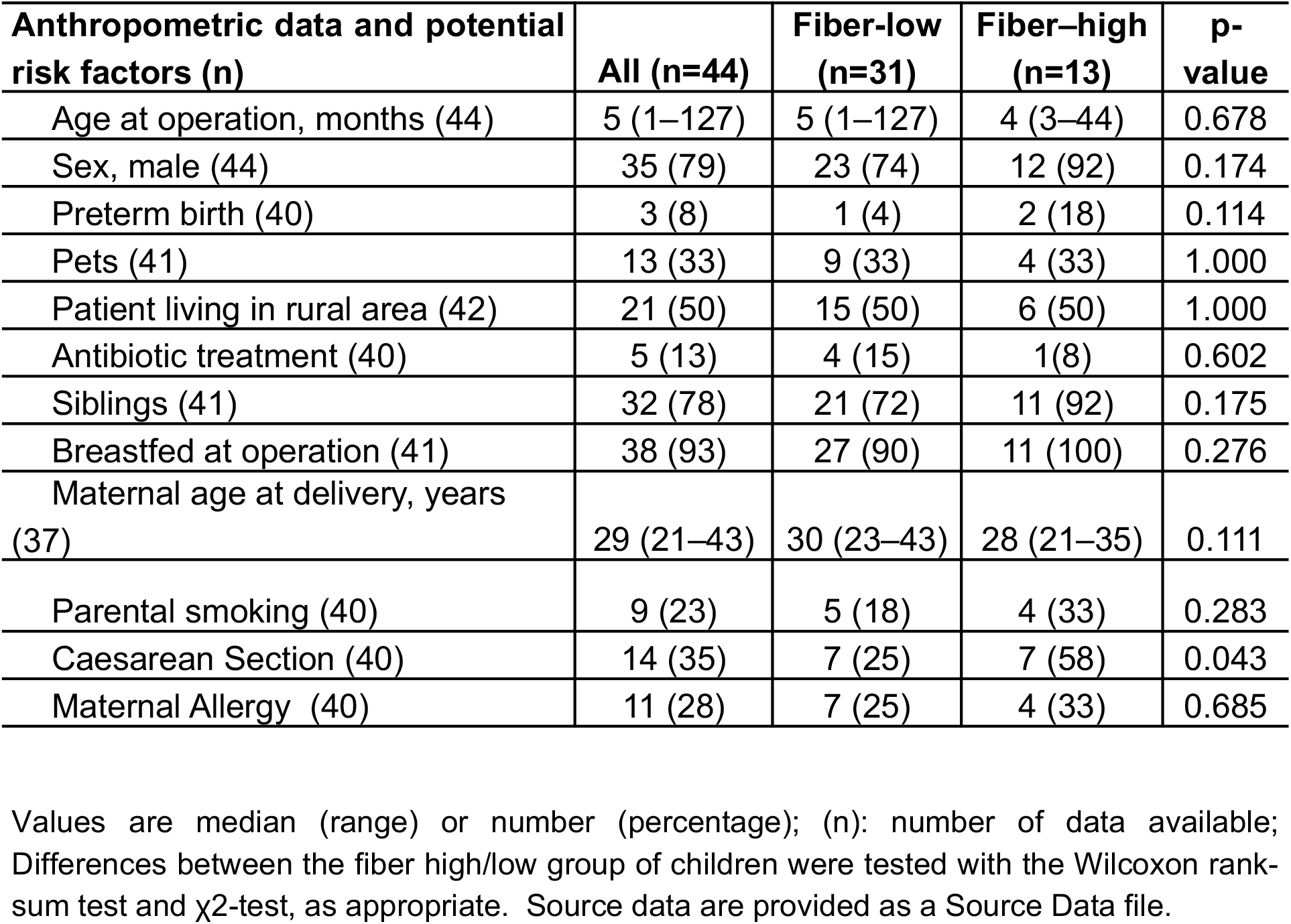
Characteristics prospective HSCR patient cohort

**Supplementary Figure 1:**
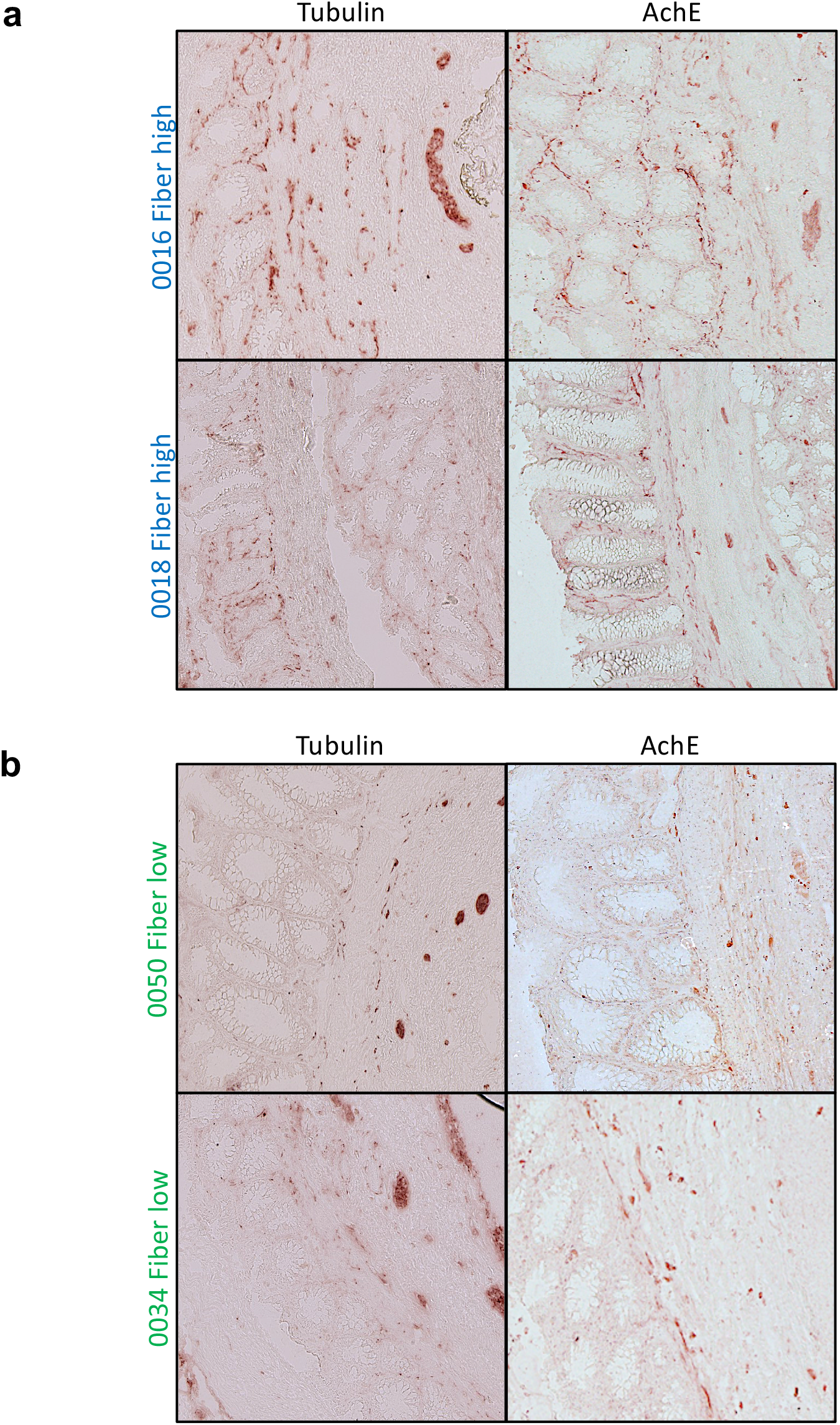
Fiber scoring of distal colon of HSCR patients using tubulin and AChE immunohistochemistry. Tubulin and AChE immunohistochemistry of 5μm cryosections of distal aganglionic colon of two fiber high (**a**) and two fiber low (**b**) HSCR patients.

**Supplementary Figure 2:**
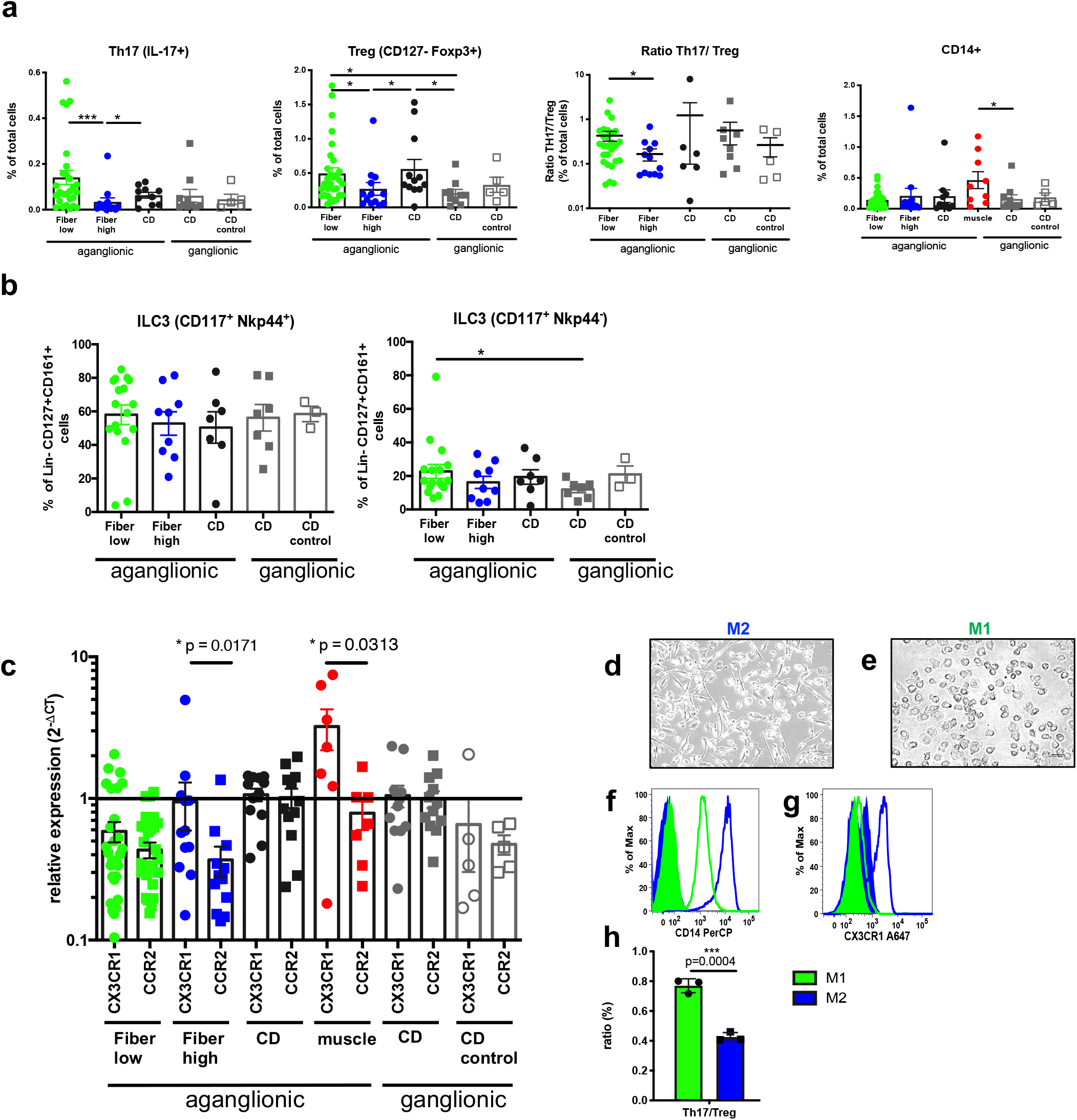
Immune cell populations in colonic tissue from HSCR and control patients and *in vitro* T-cell conversion assay. **a**, Flow cytometric analysis of mononuclear cells isolated from different aganglionic and ganglionic colonic segments of HSCR and control patients. Frequencies of total cells is shown. IL-17^+^ Th17 T cells were gated in viable CD3^+^CD4^+^ lymphocytes restimulated with PMA/Ionomycin. Treg cells were defined as CD127^-^Foxp3^+^ cells in unstimulated viable CD3^+^CD4^+^ lymphocytes. Fiber-low (n=30); fiber-high (n=13); AG CD (n=10); G CD (n=10); control CD (n=5). CD14^+^ MΦ were gated in viable HLA^+^CD64^+^ cells. Fiber-low (n=26); Fiber-high (n=13); AG CD (n=10); muscle (n=10); G CD (n=10); control CD (n=5). Exact p values indicated in Supplementary Table 4 **b** Frequencies of NCR^+^ (Nkp44^+^) and NCR^-^ (Nkp44^-^) ILC3 (CD117^+^) subsets. Fiber-low (n=17); fiber-high (n=9); AG CD (n=6); G CD (n=6); control CD (n=3). Supplementary Fig. 3c shows flow cytometric gating strategy. Scatter plots show means ± SEM. Significance was determined using unpaired nonparametric Mann-Whitney test (* p≤ 0.05; ** p≤ 0.01; *** p≤ 0.001 and **** p≤ 0.0001); Exact p values indicated in Supplementary Table 4. **c**, Quantitative RT-PCR analysis of total colonic tissue or isolated muscle tissue from different aganglionic and ganglionic colonic segments of HSCR and control patients (CX3CR1;circles and CCR2;squares). Significance was determined using Wilcoxon matched-pairs signed rank test. Fiber-low (n=31); Fiber-high (n=13); AG CD (n=12); G CD (n=11); control CD (n=5). **d-e**, Representative brightfield microscopic images (10x) of M2 (**d**) and M1(**e**) after 10 days of differentiation. Scale bar 50 μm. **f-g**, Flow cytometric analysis of blood-derived M1 and M2 macrophages from adult healthy donors. Histograms show CD14 (**f**) and CX3CR1 (**g**) expression in M1 (green) and M2 (blue) macrophages as well as in unstained controls (filled histograms). Representative analysis out of 3 independent experiments is shown. **h**, Th17 and Treg conversion assay using blood-derived M1 and M2 macrophages. Frequencies of Th17 and Treg cells were determined and Th17/Treg ratio calculated. One out of 3 independent experiments is shown. Significance was determined using unpaired t-test. Source data are provided as a Source Data file.

**Supplementary Table 4:**
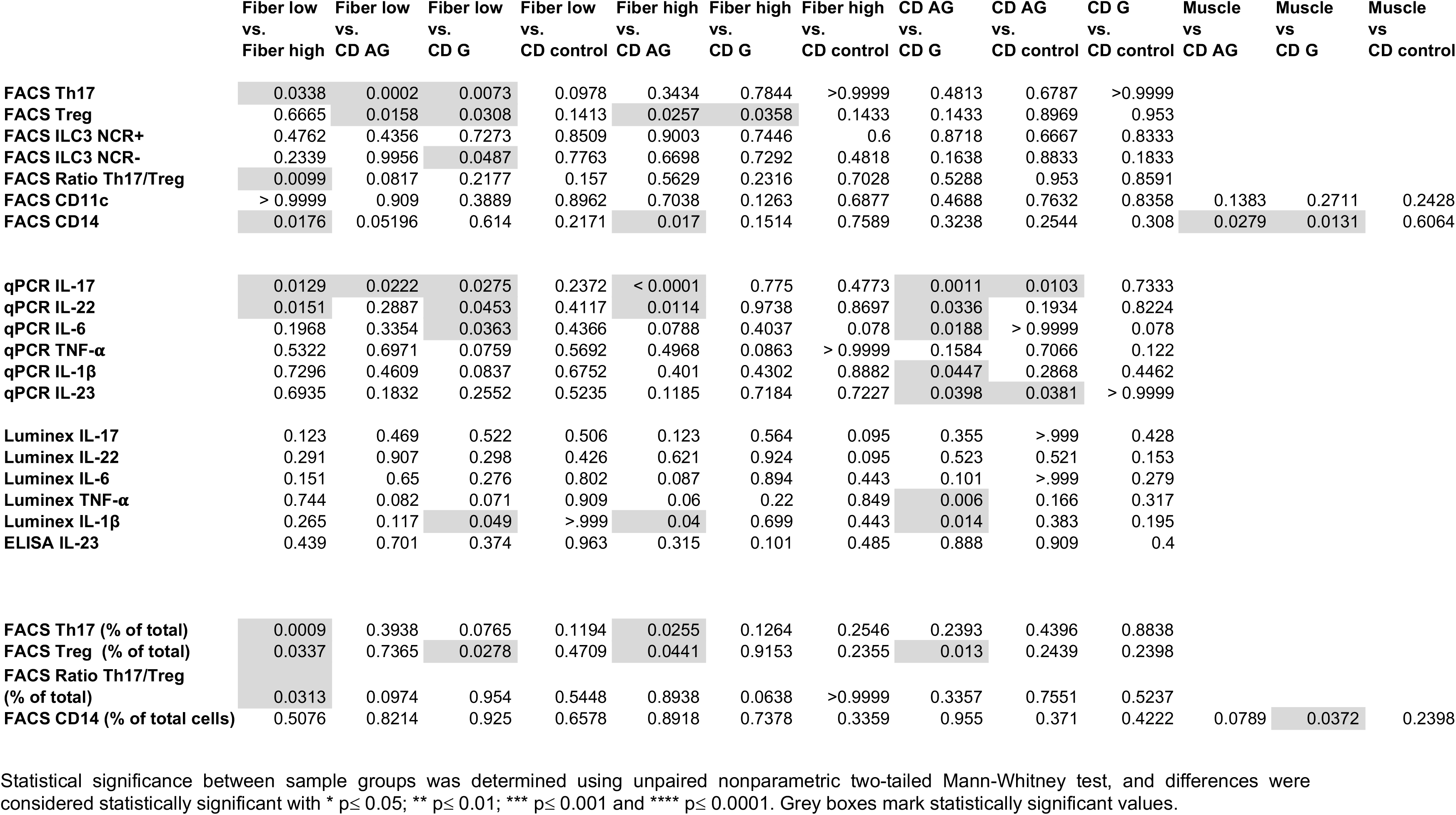
Statistical analysis of gene and protein expression

**Supplementary Figure 3:**
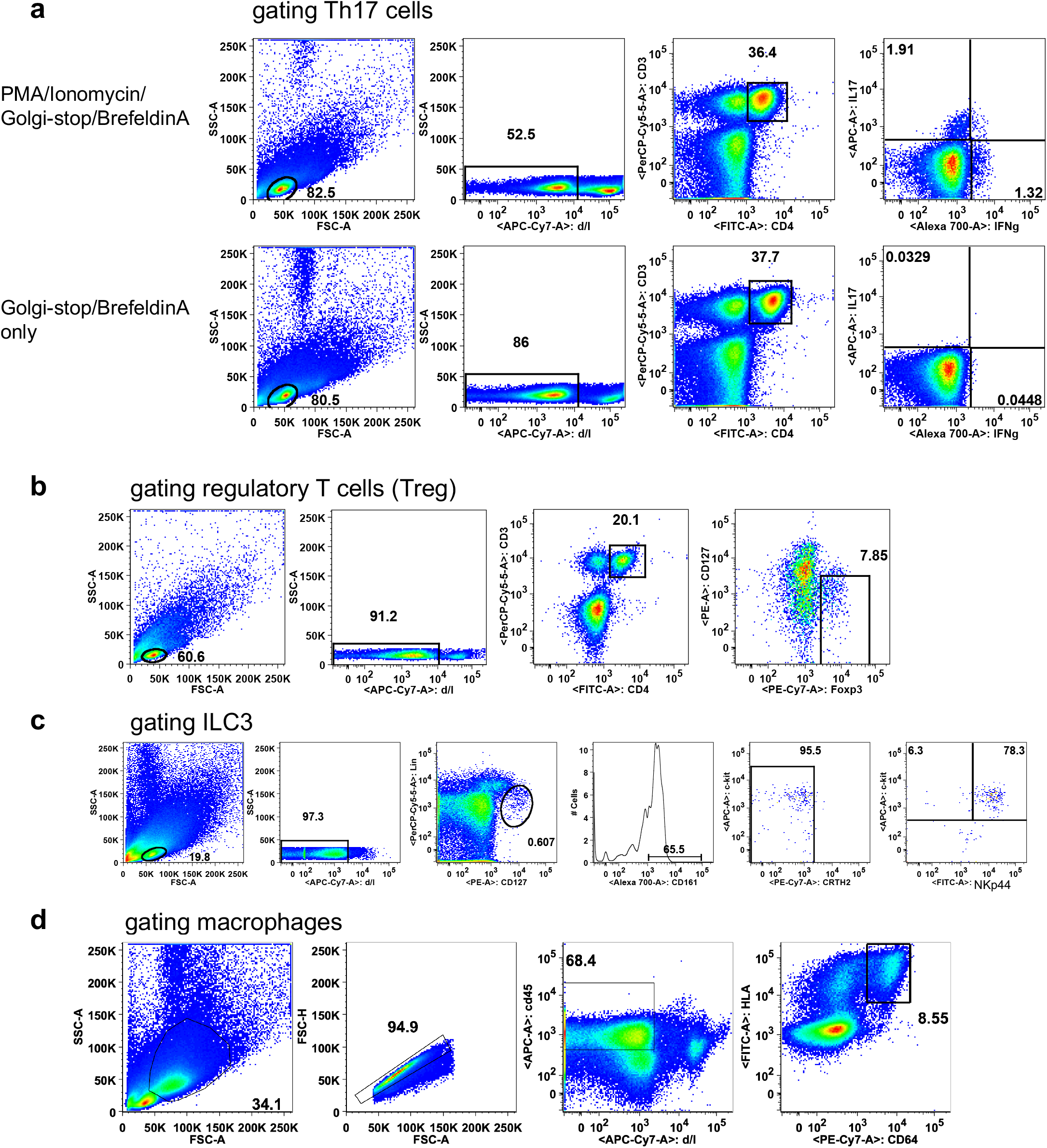
Gating strategy for flow cytometric analysis and cell sorting. **a-e**, Representative gating strategies for the FACS analysis of lamina propria mononuclear cells from HSCR and control patients. a, T helper 17 (TH17) cells were evaluated in viable CD3^+^CD4^+^ lymphocytes restimulated with PMA (phorbol 12-myristate 13-acetate)/Ionomycin in the presence of GolgiStop™ and Brefeldin A. Unstimulated Lymphocytes (Golgi-stop/Brefeldin A only) served as negative control. **b**, CD127^-^Foxp3^+^ regulatory T cells (Tregs) were gated in viable CD3^+^CD4^+^ lymphocytes. **c**, Subsets of ILC3 we**r**e detected in viable lin’CD127^+^CD161^+^CRTH2’ lymphocytes. ILC3 NCR^+^ were defined as c-kit^+^NKp46^+^ and ILC3 NCR^-^ as c-kit^+^NKp46^-^. **d**, Macrophage gating strategy for FACS analysis as well as for cell sorting. HLA^+^CD64^+^ macrophages were detected in viable CD45^+^ leucocytes.

**Supplementary Movie 1:**
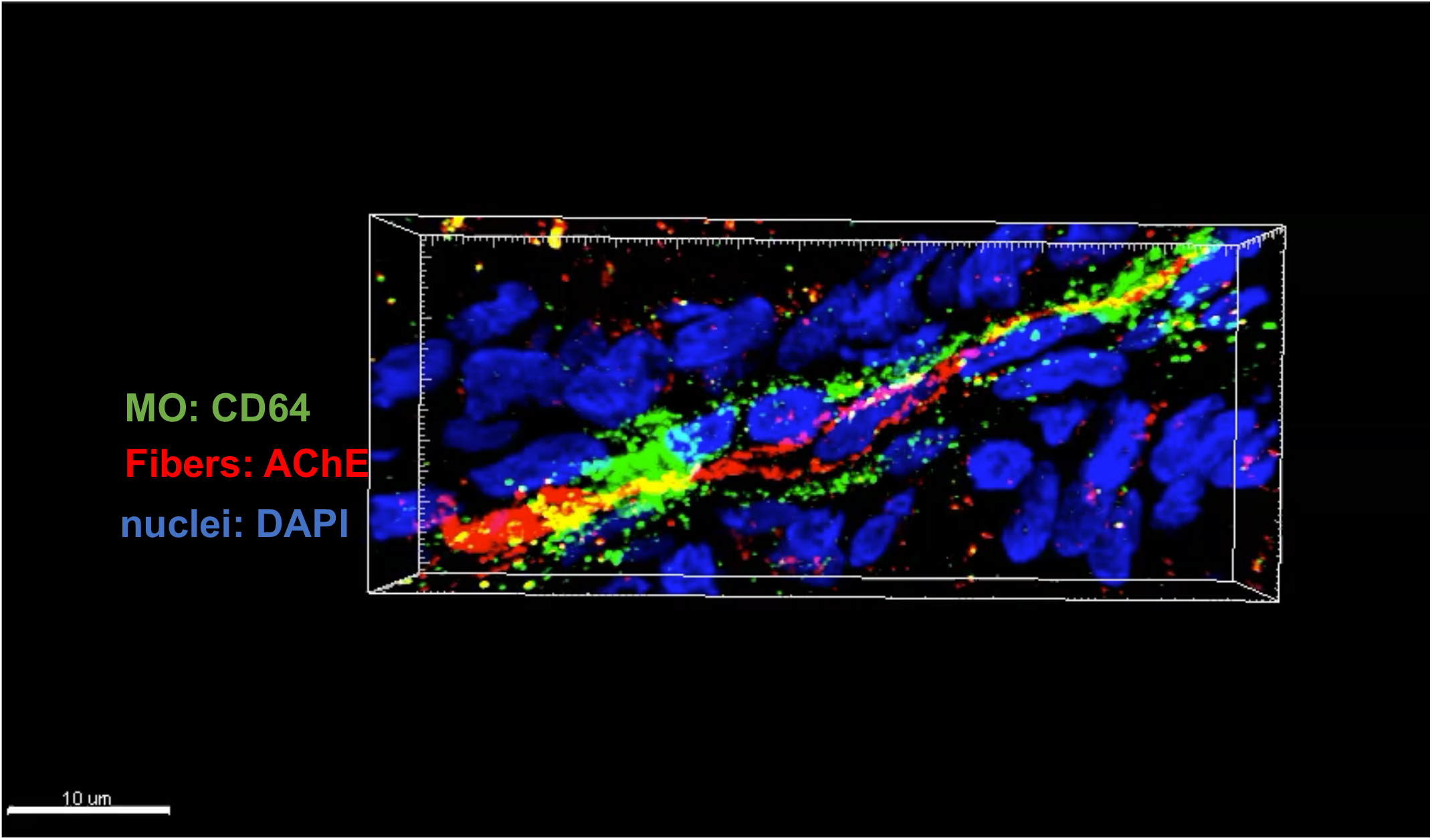
Co-localization of macrophages and cholinergic nerve fibers. Colocalization of mucosal macrophages with cholinergic nerve fibers at the epithelial region of an inter-crypt, related to Figure 4b. Cryosection (5 μm) from a rectum part of a fiber-high patient was stained using human anti-CD64 (green), anti-AChE (red), and nuclear stain (DAPI). 3D reconstruction movie from a fast scanning confocal image using Imaris software is shown.

**Supplementary Figure 4:**
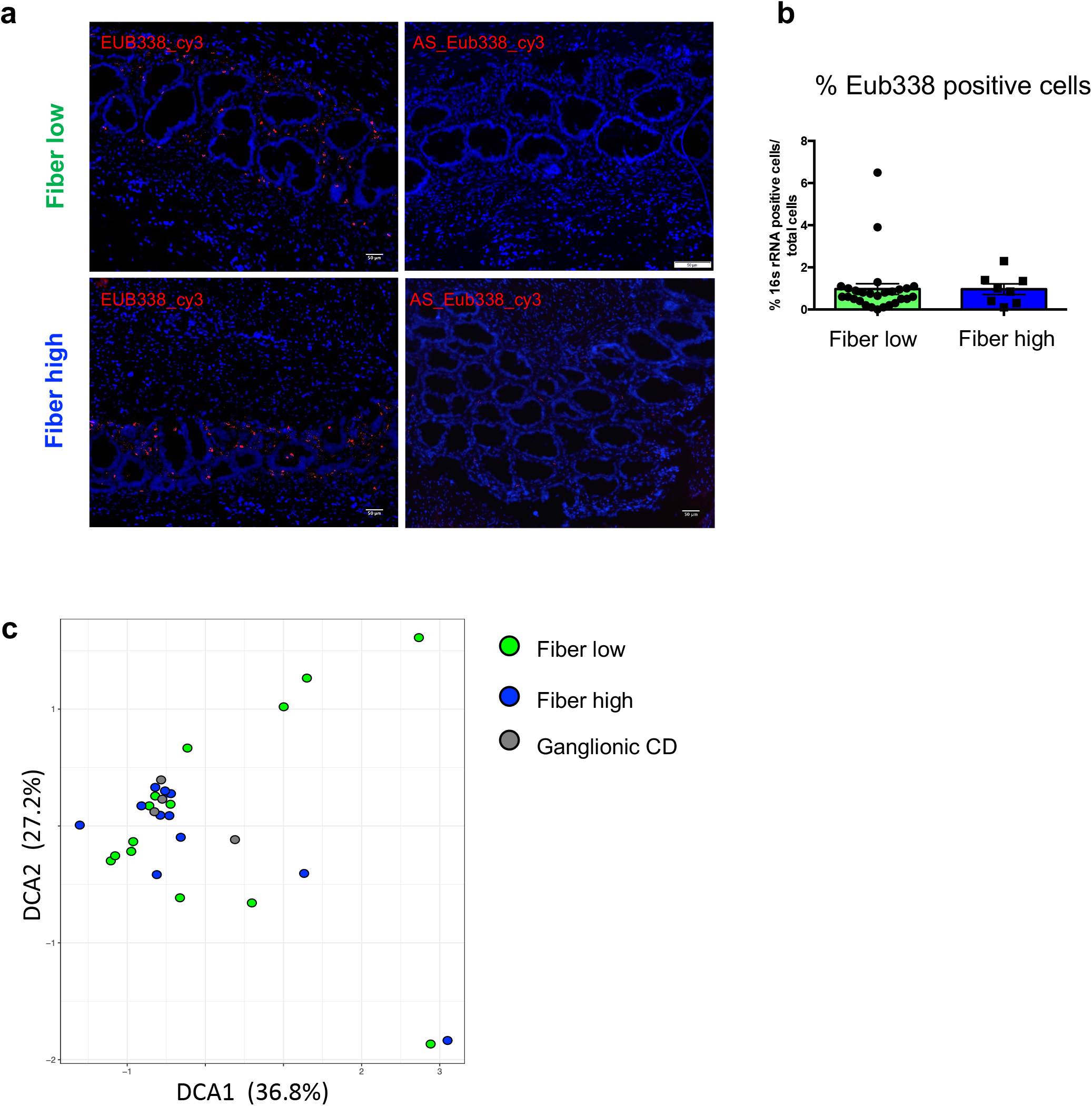
Microbial dysbiosis between fiber low and fiber high colonic tissue. **a-b**, 16s rRNA FISH analysis was performed on 5 μm cryosections of distal colon (rectum and sigmoid colon) of fiber-low (n=27) and fiber-high (n=8) HSCR patients using an EUB338-specific probe. **a**, Representative images of a fiber-low and fiber-high colonic tissue probed with Cy3-labelled sense and anti-sense (AS; negative control) EUB338. **b**, Frequencies of translocated bacteria (16sRNA^+^/DAPI^+^) were determined using CellProfiler software and are shown as percentage of total DAPI^+^ cells. **c**, 16S rRNA analysis of colonic tissue from aganglionic (fiber-low n=14; fiber-high n=11) and ganglionic (CD gang. n=4) segments of age-matched HSCR patients. Beta diversity 2D plot show patterns of intersample relations using detrended correspondence analysis (DCA). Source data are provided as a Source Data file.

**Supplementary Table 5:**
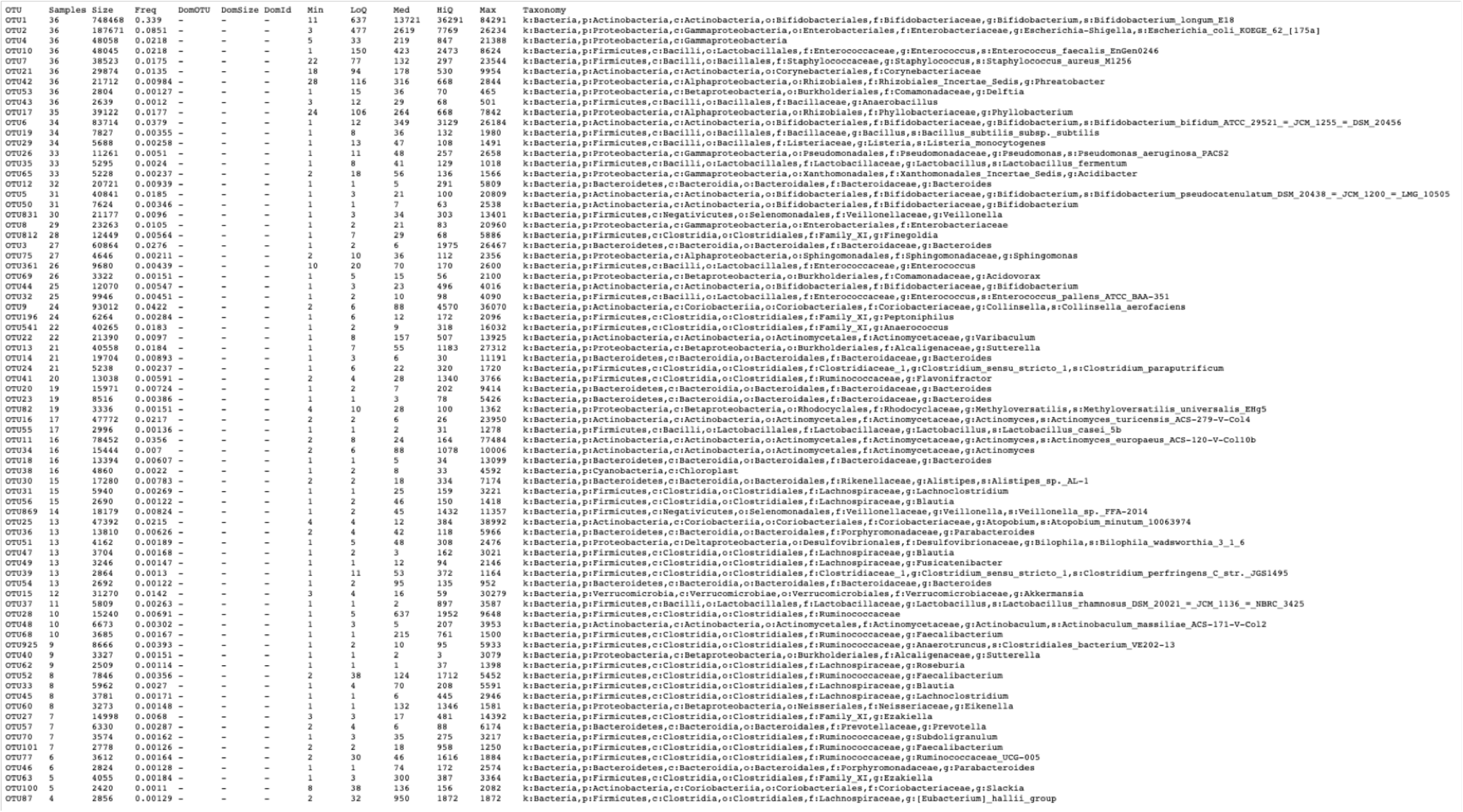
Core microbiome of 16S rDNA sequencing of colonic tissue

**Supplementary Table 6:**
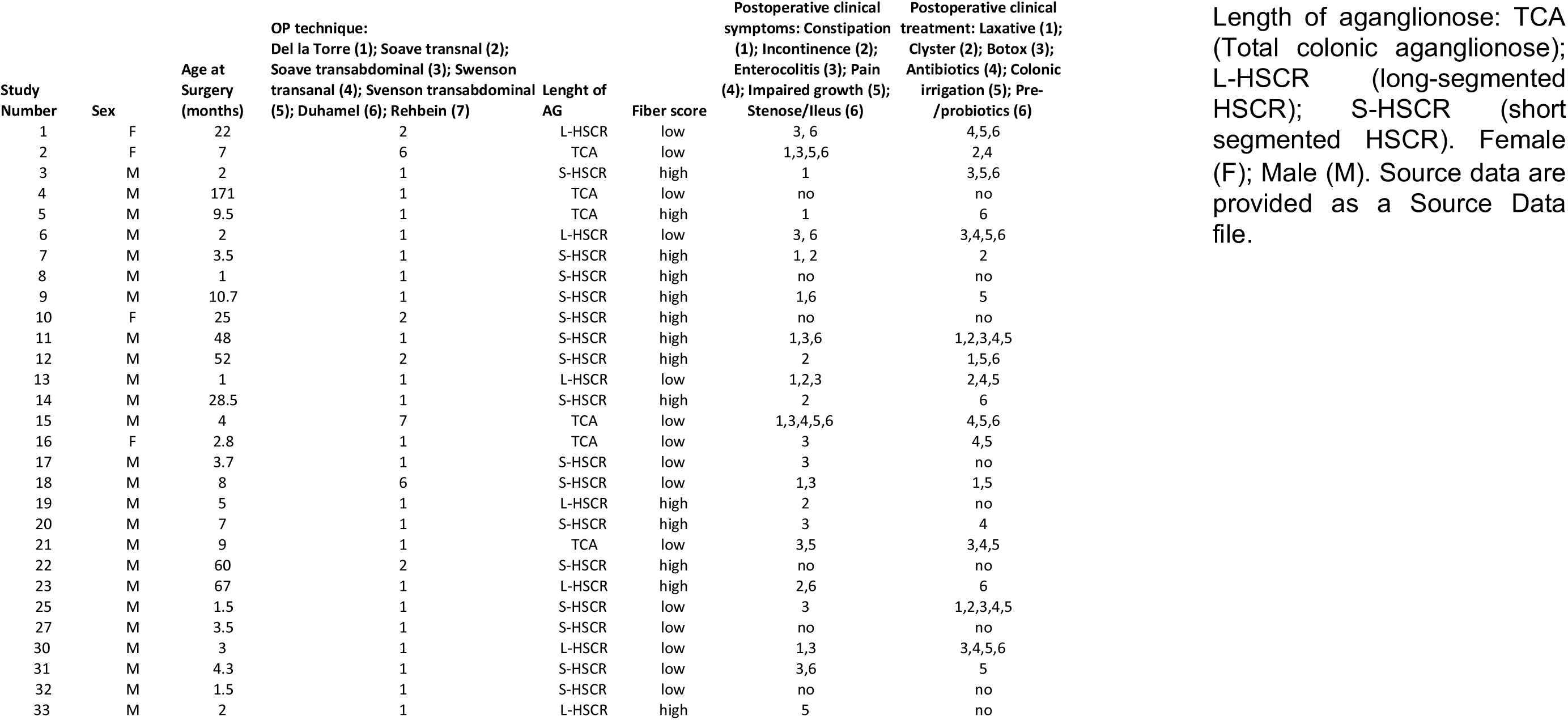
Overview of retrospective HSCR patient cohort

**Supplementary Table 7:**
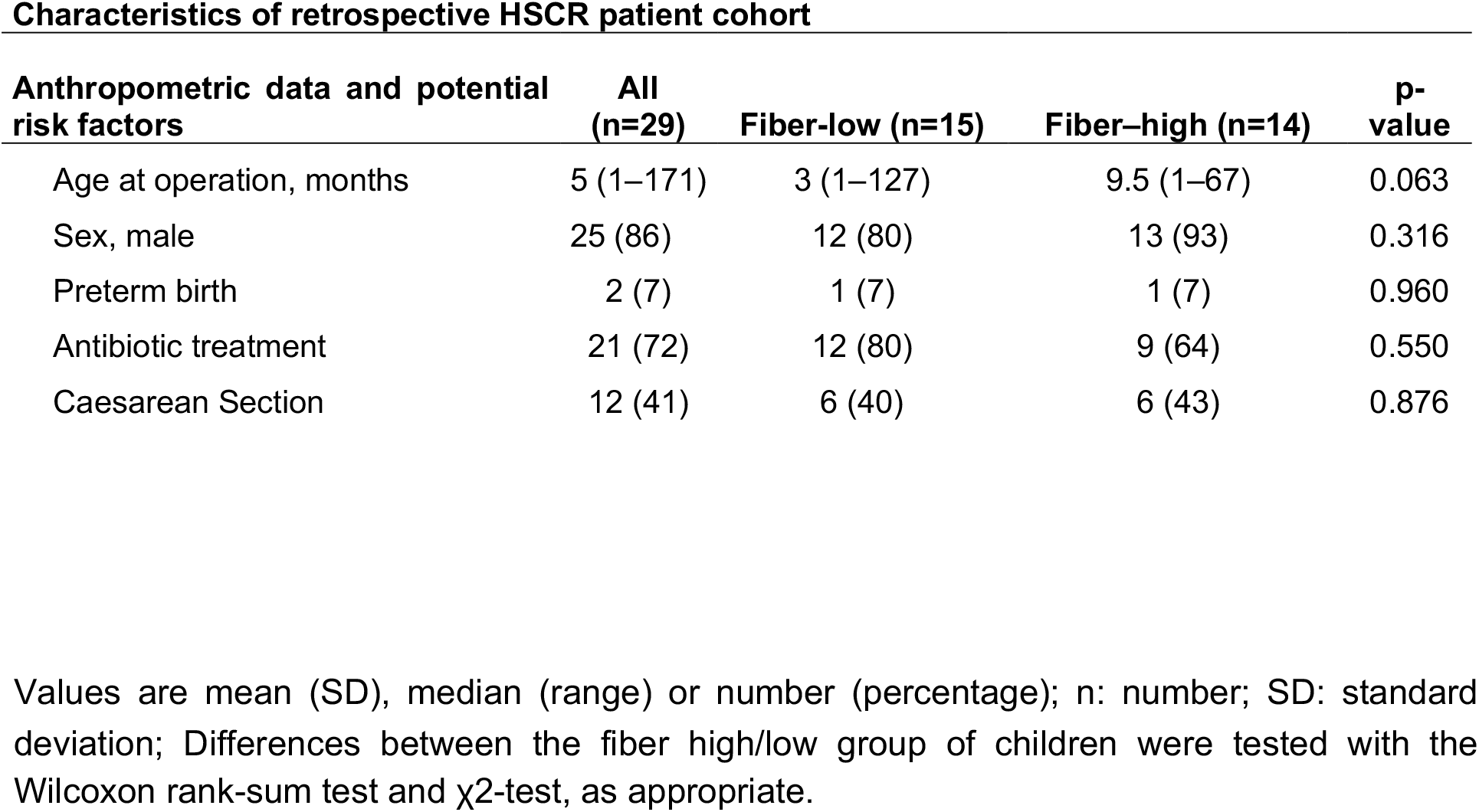
Characteristics retrospective HSCR patient cohort

**Supplementary Table 8:** Antibodies for FACS and Histology

| Antibodies for Flow cytometry |  |  |  |
| --- | --- | --- | --- |
| Name | Clone | Fluorochrome | Supplier |
| CD3 | SK7 | PerCP | BioLegend |
| HLA-DR | L243 | FITC | eBioscience |
| CD45 | HI30 | APC | BD Biosciences |
| CD14 | HCD14 | PerCP | BioLegend |
| CD64 | 10.1 | PE-Cy7 | BioLegend |
| CD11c | 3.9 | Alexa700 | eBioscience |
| CD4 | RPA-T4 | FITC | BD Biosciences |
| CD127 | A019D5 | PE | BioLegend |
| CD25 | BC96 | Alexa 647 | BioLegend |
| NKp44 | p44-8 | BB515 | BD Biosciences |
| CD11c | 3.9 | PerCP/Cy5.5 | BioLegend |
| CD19 | HIB19 | PerCP | BioLegend |
| CD56 | HCD56 | PerCP | BioLegend |
| FcεRI | AER-37 | PerCP | Novus Biologicals |
| TCR α/β | IP26 | PerCP/Cy5.5 | BioLegend |
| TCR γ/δ | B1 | PerCP/Cy5.5 | BioLegend |
| CD117 (c-kit) | 104D2 | APC | BioLegend |
| CD294 (CRTH2) | BM16 | PE-Cy7 | BioLegend |
| CD161 | HP-3G10 | Alexa 700 | BioLegend |
| IL-17A | eBio64DEC17 | APC | eBioscience |
| IFN-γ | B27 | Alexa 700 | BioLegend |
| Roryt | Q21-559 | Alexa 488 | BD Biosciences |
| Foxp3 | PCH101 | PE-Cy7 | eBioscience |
| CD16/CD32 (Fc block) |  | purified | BD Biosciences |

| Primary Histology antibodies |  |  |  |  |
| --- | --- | --- | --- | --- |
| Name | clone | fluorochrome | host/subtype | supplier |
| CD64 | 10.1 | purified | mouse IgG1 | BioLegend |
| CD3 | UCHT1 | Alexa 488 | mouse IgG1 | BioLegend |
| Acetylcholinesterase (AchE) | HR2 | purified | mouse IgG2b | Abcam |
| Beta III tubulin | 2G10 | purified | mouse IgG2a | Abcam |
| Beta III tubulin | GT11710 | purified | mouse IgG2b | Genetex |
| Tyrosine Hydroxylase (TH) | 1A5 | purified | mouse IgG2a | Invitrogen |
| nNOS | 16/nNOS/NOS type | purified | mouse IgG2a | BD Bioscience |
| S100b | 11C12E12 | purified | mouse IgG1 | Biolegend |
| VIP | polyclonal | purified | guinea pig | Peninsula Laboratories |
| IL-23p19 | 23dcdp | PE | mouse IgG2b | eBioscience |

| Secondary Histology antibodies |  |  |  |  |
| --- | --- | --- | --- | --- |
| Name | clone | fluorochrome | host/subtype | supplier |
| anti-mouse IgG1 | polyclonal | Alexa 488 | goat | Invitrogen |
| anti-mouse IgG1 | polyclonal | Alexa 555 | goat | Invitrogen |
| anti-mouse IgG1 | polyclonal | Alexa 647 | goat | Invitrogen |
| anti-mouse IgG2a | polyclonal | Alexa 647 | goat | Invitrogen |
| anti-mouse IgG2a | polyclonal | Alexa 555 | goat | Invitrogen |
| anti-mouse IgG2b | polyclonal | Biotin | goat | BD Biosciences |
| anti-mouse IgG2b | polyclonal | Alexa 647 | goat | Invitrogen |
| anti-guinea pig | polyclonal | Alexa 555 | goat | Invitrogen |
| Streptavidin |  | Cy3 |  | BioLegend |

**Supplementary Table 9:**
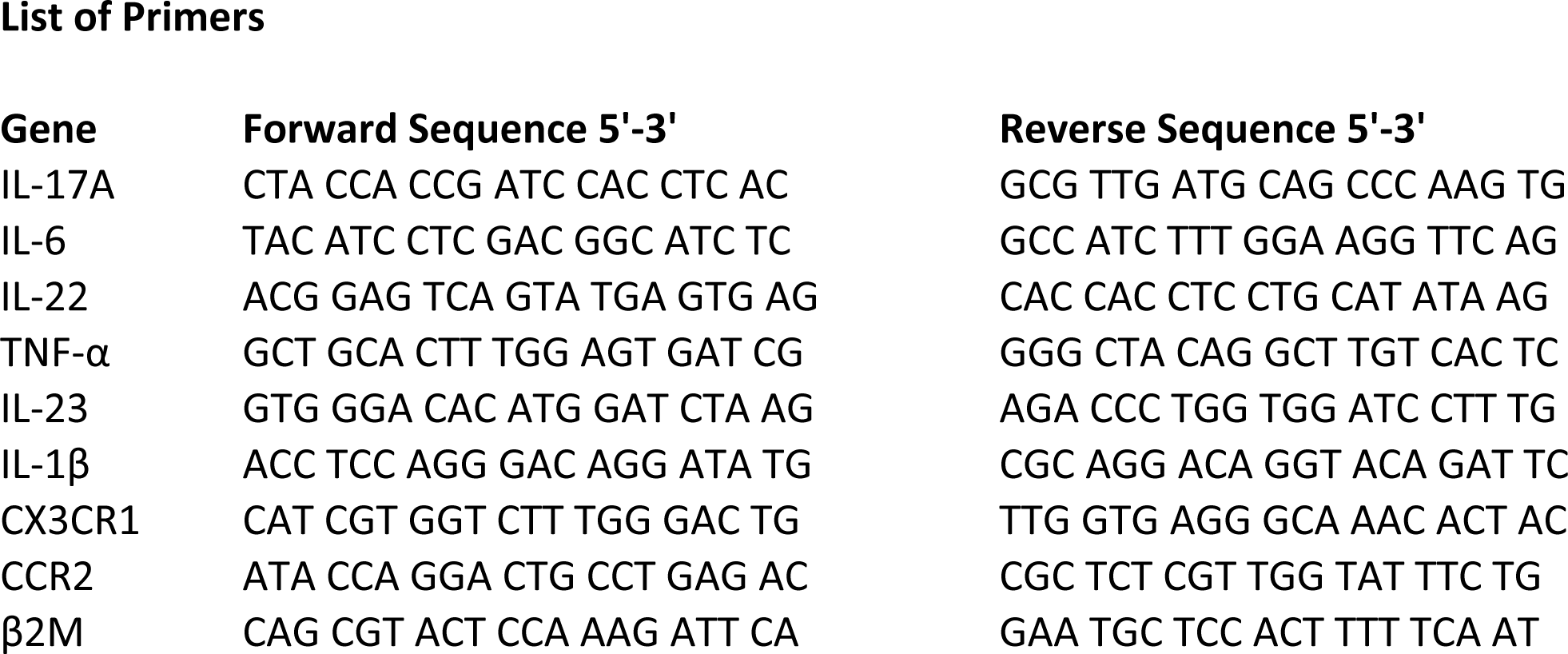
Primers

**Supplementary Figure 5:**
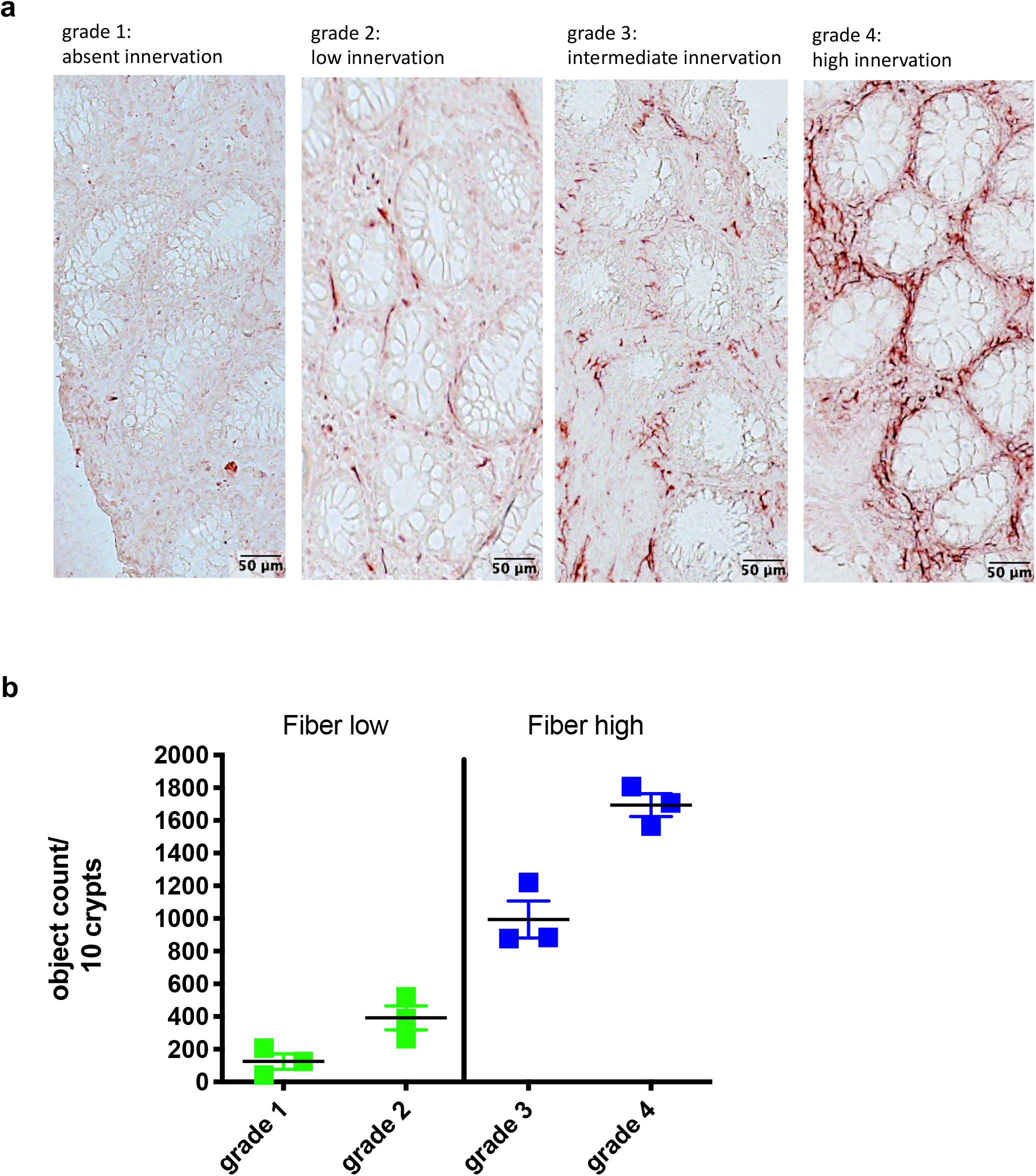
Fiber innervations grades for Fiber scoring. **a**, AChE immunohistochemistry of 5μm cryosections of distal aganglionic colon showing 4 different mucosal innervation grads. Grades 1 and 2 were grouped into fiber-low and 3 and 4 were grouped into fiber-high HSCR patients. **b**, Using brightfield microscopy and CellSens Dimension Software the innervation grade was quantitatively confirmed in 3 representative patients per group (for details see methods). Scatter dot plots show means ± SEM. Source data are provided as a Source Data file.

