## Supplementary material for "Lack of mucosal cholinergic innervation is associated with increased risk of enterocolitis in Hirschsprung’s disease": Methods

### Supplementary Materials and Methods

#### Patients and human specimens

A total of 44 children (median age 5 months) with HSCR as well as miscellaneous intestinal diseases (non-HSCR, control group) were enrolled in a prospective multicenter study conducted in Switzerland and Germany. The study was approved by the Ethics Committees of Northwest/Central Switzerland (EKNZ 2015-049) and the Medical Faculty Heidelberg (S-388/2015). The study was registered at www.clinicaltrials.gov (accessing number NCT03617640) and was performed according to the Declaration of Helsinki. After obtaining consent, colonic tissue was collected freshly in the operating room. For histological analysis, a longitudinal strip of resected colonic tissue was immediately snap-frozen. Tissue for RNA analysis was stored in RNAlater (Qiagen) or immediately frozen on dry ice. Tissue for fluorescence-activated cell sorting (FACS) analysis was kept on ice in Hanks' Balanced Salt Solution (HBSS) supplemented with 100 units penicillin, 0.1 mg/mL streptomycin, 0.25 µg/mL amphotericin B, and 40 µg/mL gentamycin and was processed in less than 24 h. The HSCR group included patients with short-segment (n=31), long-segment (n=6), and total (n=7) colonic aganglionosis. Surgery was performed according to the Swenson or Soave technics (Supplementary Table 1)[114-116]. The length of aganglionosis was determined intraoperatively in mapping biopsies, and postoperatively in the resected tissue via enzyme-histochemical staining. Tissue staining and assessment were performed by the pathology units of the respective clinics (Supplementary Table 2). Ganglionic tissue was characterized by the presence of intrinsic nerve cell bodies. The presence of intrinsic nerve cell bodies (ganglia) was detected by several enzymatic stainings: lactic dehydrogenase (LDH), succinic dehydrogenase (SDH), nitroxide synthase (NOS), nicotinamide adenine dinucleotide phosphate (NADPH), and immunohistochemical calretinin staining (Supplementary Table 2). Aganglionic tissue was characterized by the lack of intrinsic cell bodies and presence of extrinsic nerve fibers in the distal colon. Extrinsic nerve fibers were detected by enzymatic acetylcholinesterase (AChE) staining (Supplementary Table 2)[16,33]. Study group characteristics (clinical and nonclinical variables) were evaluated via a parent questionnaire. Postoperative symptoms (obstipation, incontinence, enterocolitis, and pain) as well as postoperative treatments (laxative, clyster, botox, antibiotics, colonic irrigation, and pre-/probiotics) were evaluated from medical records one year after surgery. Enterocolitis was defined to fulfill at least 3 of the following criteria: diarrhea (>3x/day), vomiting, temperature >38.5°C, antibiotics, hospitalization, meteorism, and high inflammatory blood parameters (i.e., leucocytes, C-reactive protein (CRP)). For the retrospective cohort analysis, we included HSCR patients diagnosed and treated at the University Children's Hospital Basel between 2003 and 2018. Patients were identified using International Classification of Disease (ICD-10) Diagnosis Q43.1. Swiss roll cryosections (5 µm) from resected distal aganglionic colon were provided by the Department of Pathology, University Hospital Basel, Basel, Switzerland. Fiber scoring was performed using beta III tubulin immunohistochemistry and evaluated in a double-blind fashion by three individuals. Demographic and clinical data were extracted from medical records. Enterocolitis was defined in the same way as in the prospective cohort study. The study was approved by the Ethics Committee of North-West-Switzerland (EKBB 2019-01406).

**Immunohistochemistry and nerve fiber scoring**

Longitudinal colonic strips were washed in ice-cold phosphate buffered saline (PBS) and embedded as a Swissroll in Tissue-Tek® O.C.T. Compound (Sakura).

Cholinergic nerve fibers were visualized in 5 µm cryosections using mouse IgG2b anti-human AChE (Abcam) or mouse IgG2a anti-human beta III tubulin antibody together with an anti-mouse HRP-AEC staining kit (R&D) according to instruction manuals. Slides were automatically scanned using an Olympus BX63 motorized brightfield microscope using a 10x objective. Presence of extrinsic nerve fibers in mucosal regions of the distal aganglionic colon was evaluated semi-quantitatively in a double-blind investigation by four individuals. Mucosal regions were classified into 4 different fiber innervation grades according to absent, low, intermediate, and high innervation density (Supplementary Fig. 5a). Innervation grade 1 and 2 were grouped into fiber-low and 3 and 4 were grouped into fiber-high HSCR patients. For the quantitative measure of the $AChE^{+}$ nerve fibers in the mucosal regions images were analyzed using brightfield microscope and CellSens Dimension 2.2 Software (Olympus). In average 6 cropped region of interest containing 10 epithelial crypts from distal colon, resulting in 60 crypts per patients, were analyzed. In total 12 patients (3 patients per innervation grade) were chosen according to the visual scoring. A defined manual threshold was applied on each cropped image and object counts were obtained by the Olympus CellSens dimension software (Supplementary Fig. 5a).

**Isolation of mononuclear cells from colonic tissue**

Colonic specimens were separated into rectum, sigmoid colon, colon descendens, transverse colon, and colon ascendens (distal to proximal) according to the length of the resected colon[117]. Excess fat was removed, and muscle layers were stripped from the mucosa. Mucosa and muscle tissue were minced and digested separately with collagenase D (1 mg/mL) and DNase I (4 µg/mL, both Roche Diagnostics, Basel?) in complete medium (RPMI-1640 supplemented with 10% FCS, 200 units penicillin, 0.2 mg/mL streptomycin, 0.5 µg/mL amphotericin B, 80 µg/mL gentamycin, and 10 mM Hepes) at 37°C for 1 h under vigorous shaking. Digestion suspension was filtered through a 100 µm cell strainer, and cells were washed in complete medium. Suspension cells were resuspended in 20% percoll (3 mL) and underlaid with 40% percoll and 70% percoll (3 mL of each). Mononuclear cells were purified from the 40/70% percoll interface whereas epithelial cells were excluded in the 20/40% percoll interface. Cells were used freshly or were cryopreserved for further analysis.

**Flow cytometry and fluorescence-activated cell sorting (FACS)**

For intracellular cytokine detection, cells were restimulated in complete medium with phorbol 12-myristate 13-acetate (PMA; 150 ng/mL, Sigma-Aldrich) and ionomycin (750 ng/mL, Sigma-Aldrich) in the presence of GolgiStop$^{TM}$ and Brefeldin A (BD Biosciences) for 4 h at 37°C in a humidifying atmosphere. Viability was determined using LIVE/DEAD Fixable Near-IR dead cell stain (ThermoFisher). Cell surface markers were stained extracellularly (15 min, 4°C) in FACS buffer (PBS supplemented with 2% fetal calf serum, FCS), whereas cytokines and transcription factors were detected intracellularly using Cytofix/Cytoperm (BD Biosciences) or Foxp3/transcription factor staining buffer set (eBioscience), respectively. Unspecific Fcγ receptor binding was blocked by human Fc block (CD16/CD32). Lineage cocktail for innate lymphoid cells (ILC) contained markers for T cells (CD3, TCRα/β; TCRγ/δ), NK T cells (CD56), B cells (CD19), dendritic cells, monocytes, MΦ (CD11c, CD14, CD16), and mast cells (FcεRI). Cells were analyzed using FACS Canto II (BD Biosciences) and FlowJo software (TreeStar Inc.). MΦ were sorted as viable $CD45^{+}HLA^{+}CD64^{+}$ using FACS Aria cell sorter flow cytometry (BD Biosciences). Supplementary Table 8 details the human specific antibodies.

**RNA isolation and quantitative real-time PCR (qRT-PCR)**

Colonic tissues samples (max. 30 mg) were homogenized in RLT lysis buffer containing 1% 2-mercaptoethanol (Sigma-Aldrich) using a tissue disruptor (IKA). Total RNA from tissue or cells was isolated using the RNAeasy Plus mini kit (QiAgen) and was reverse-transcribed using GoScript Reverse Transcription System (Promega). Subsequently, qRT-PCR was done using the ViiA7 RT-PCR system (Thermo Fisher Scientific) and FastStart Universal SYBR Green Master (Roche Diagnostics) according to the manufacturers' instructions. Relative gene expression was calculated using the $2_{-\Delta CT}$ method, with β2-microglobulin as housekeeping gene, and results were multiplied by a factor of 1000. Primer pairs were designed according to exon junction span using the clone manager software (Sci-Ed Software). Supplementary Table 9 details the human-specific primer pairs.

**Generation of blood-derived M1 and M2 MΦ**

Peripheral blood mononuclear cells (PBMCs) from healthy adult volunteers were isolated using the SepMate/Lymphoprep system (Stemcell) according to the user manual. For monocyte attachment, PBMCs were resuspended in RPMI (supplemented with 10% FCS, 200 units penicillin, and 0.2 mg/mL streptomycin) and seeded into T-75 flasks for 2 h at 37°C in a humidifying atmosphere. The attached monocytes were washed twice with PBS and then cultured for 10 days in M1 or M2 Macrophage Generation Medium DXF (Promocell) according to the manufacturer's instructions. On day 10, cells were detached by adding Accutase® cell detachment solution (Sigma-Aldrich), washed twice with RPMI, and used for further applications.

**T cell conversion assay**

Autologous naïve CD4 T cells were isolated from PBMCs using EasyStep Human naïve $CD4^+$ T cell Isolation Kit (Stemcell) according to instruction manuals. M1 or M2 MΦ ($10^4$/well/200 µL) were cocultured with autologous naive $CD4^+$ T cells ($10^5$/well/200 µL) in the presence of 5 µg/mL anti-human CD3 for 6 days at 37°C in a humidifying atmosphere. Alternatively, $CD4^+$ T cells were activated using anti-human CD3/CD28 activation Dynabeads (Life Technologies) in a 1:1 ratio. For T helper cell differentiation, we added the following cytokines: IL-2 (5 ng/mL) and TGF-β1 (2 ng/mL) for Treg conversion and TGF-β1 (2 ng/mL) and IL-6 (10 ng/mL) for Th17 differentiation. All recombinant human cytokines were purchased from Preprotech. On day 6, T cells were separated from MΦ or Dynabeads into a new well and restimulated with PMA (150 ng/mL) and ionomycin (750 ng/mL) in the presence of GolgiStop$^{TM}$ and Brefeldin A for 4 h at 37°C in a humidifying atmosphere. T cell subsets were subsequently analyzed by flow cytometry using fluorescent antibodies against human TCRα/β, CD4, and IL-17/Rorγt or CD25/Foxp3 for Th17 and Treg cell identification. Cytokines and transcription factors were detected intracellularly using transcription factor buffer set (BD Biosciences) according to the manufacturer's instructions. Frequencies of single subsets were determined as percentages of viable CD4/TCRα/β positive T cells. Ratios of Th17/Treg frequencies were calculated.

**Immunofluorescence and colocalization studies**

Longitudinal colonic strips were washed in ice-cold PBS and embedded as a Swiss roll in Tissue-Tek® O.C.T. compound. For fluorescence staining of immune cells and cholinergic fibers, we established a modified protocol of the anti-mouse HRP-AEC staining kit (R&D). All steps were performed at room temperature. Between all steps, the tissues were washed three times for 10 min in PBS and antibodies/streptavidin/4’,6-diamidine-2’-phenylindole dihydrochloride (DAPI) were diluted in antibody diluent. Briefly, cryosections (5 µm) were fixed with 4% paraformaldehyde (PFA) for 5 min, followed by a blocking step with serum blocking reagent (15 min). Subsequently avidin and biotin were blocked using Avidin Blocking Reagent (15 min) followed by Biotin Blocking Reagent (15 min). Primary antibodies, purified mouse IgG2b anti-human AChE, together with purified mouse IgG1 anti-human CD64 or mouse IgG1 anti-human CD3-A488 or mouse IgG2a anti-human β3tubulin, were incubated for 1 h. CD64, AChE, β3tubulin, TH, nNOS, S100b, and VIP were detected by secondary antibodies goat anti-mouse IgG1 A488, goat anti-mouse IgG2b biotin, goat anti-mouse IgG2a A647 or IgG2b A647, goat anti-mouse IgG2a A555, goat anti-mouse IgG2a A555, goat anti-mouse IgG1 A555, and goat anti-guinea pig A555, respectively. IL-23p19 was directly PE-labeled. Biotin was visualized by streptavidin-Cy3 incubation for 30 min. Finally, nuclei were visualized by DAPI staining (3 min; 0.5 µg/mL), and slides were mounted using Prolong Diamond Antifade Mountant (Life Technologies). Secondary antibody controls were included as negative controls. Images were taken by a fluorescence microscope (BX43 Olympus) and analyzed using CellSense (Olympus) and Fiji. Supplementary Table 8 details the antibodies used for histology. Fast-scanning confocal microscopy was performed with a Zeiss LSM880 Airyscan inverted system. Images were acquired and processed using Zen Black with a 63x plan apochromatic oil immersion objective DIC M27(NA1.4) using the three respective Ar-lasers 561 nm, 488 nm, and 405 nm in 4 frames average fast mode. The acquisition parameters and settings of the images were verified for the absence of crosstalk between Green and Red excitation and gain of the different lasers was always set at the same values for all acquisitions. Negative staining controls (secondary antibodies only) were used to determine pixel intensity background for the quantification. Images of fiber-high and fiber-low tissues were used for acquisition. Using a Fiji software macro, we quantified the colocalization pixel intensity between MΦ (or T cells) and AChE fibers. Briefly, images were created by multiplying and subsequent automated thresholding of the respective channels. Regions of interest containing MΦ (or T cells) were manually annotated in the raw data. The sum of colocalized pixel values contained within 100 to 200 annotated MΦ (or T cells) per patient tissue were measured on a sum projection image. Data are shown as raw integrated density (RawIntDen).

For the quantification of IL-23, images recorded under a 20x objective were analyzed using CellSense Software (Olympus) and Cellprofiler-3.1.9. To quantify IL-$23^+$ MΦ in tissue of different patient groups (4 fiber-low, 4 fiber-high), 4 to 8 regions per patient from rectum and sigmoid colon were used. Primary objects were defined by nuclear DAPI staining. Secondary object was defined by extracellular CD64 expression using the watershed module after applying a global threshold for improved cell identification. IL23 positive cells were identified as a second primary object ($IL23^+$), using Three classes Otsu thresholding method, and subsequently related to CD64 MΦ. Frequencies of IL-$23^+$ MΦ ($IL23^+CD64^+$) were shown as the percentage of total MΦ.

**16s rRNA Fluorescence in situ hybridization**

The FISH procedure was performed in 5 µm rectum/sigmoid colon Swiss roll cryosections. Colonic tissue sections were fixed with 4% PFA for 20 min on ice followed by a wash step in PBS. Slides were dipped into ice-cold methanol prior to hybridization with Cy3-labeled EUB338 sense probe (GCT GCC TCC CGT AGG AGT) or EUB338 anti-sense probe (ACT CCT ACG GGA GGC AGC) as negative control, in hybridization buffer (20 mM Tris-HCl; 0.9% NaCl; 0.1% SDS) at 48°C for

16 h. Slides were washed twice in hybridization wash buffer (20 mM Tris-HCl; 0.9% NaCl; pH 7.2) for 30 min at 48°C and subsequently mounted with ProLong Gold Antifade Mountant with DAPI (Life Technologies) and evaluated with a fluorescence microscope (BX43 Olympus). Images were analyzed using CellSense Software (Olympus) and Fiji software. To quantify the bacterial translocation in the different patient groups, 4 to 6 regions per picture from rectum and sigmoid colon were used, and corresponding sites were used for anti-sense probe. On average, 2000 DAPI$^{+}$ cells/region were then processed using Cellprofiler software. In brief, primary objects were defined by nuclear DAPI staining. After masking the nuclei, a second primary object (Eub338_bacteria) based on nuclear Eub338 expression was defined and related to DAPI$^{+}$ cells. Global Otsu two classes thresholding method was applied for DAPI nuclei staining, and manual thresholding for Eub338 was used before masking the nuclei to subtract the background. The anti-sense probe was running in parallel under the same pipeline, and results were subtracted from the sense probe. Frequencies of translocated bacteria (16sRNA$^{+}$/DAPI$^{+}$) were expressed as the percentage of total DAPI$^{+}$ cells.

**16S rDNA gene sequencing**

For 16s rDNA microbial analysis, we used colonic cryosections (100 µm). Extraction, lysis, and DNA isolation were done using Fast DNA Stool Mini Kit (Qiagen) according to the manufacturer's instructions. Bead beating was run on a fastprep24 instrument (MPBiomedicals; 2 cycles of 45 s at speed 4, followed by 2 cycles of 45 s at speed 5) in 2 mL screwcap tubes containing 0.1 mm zirconia beads (0.3 g) and 2.3 mm zirconia beads (0.36 g). The whole raw extract was prepared for DNA isolation. Concentration of the isolated DNA was determined with PicoGreen measurement (Quant-iT™ PicoGreen™ dsDNA Assay Kit, Thermo Fisher), and integrity was checked by agarose gel electrophoresis.

The Illumina MiSeq platform and a v2 500 cycles kit were used to sequence the PCR libraries. The produced paired-end reads which passed Illumina's chastity filter were subjected to de-multiplexing and trimming of Illumina adaptor residuals using Illumina's real time analysis software included in the MiSeq reporter software v2.6 (no further refinement or selection). The quality of the reads was checked with the software FastQC version 0.11.7. The locus-specific V34 primers were trimmed from the sequencing reads with the software Cutadapt v1.14. Paired-end reads were discarded if the primer could not be trimmed. Trimmed-forward and reverse reads of each paired-end read were merged to reform the sequenced molecule *in silico*, considering a minimum overlap of 15 bases using the software USEARCH version 10.0.240. Merged sequences were then quality-filtered, allowing a maximum of one expected error per merged read and also discarding those containing ambiguous bases. The remaining reads were denoised using the UNOISE algorithm implemented in USEARCH to form operational taxonomic units (OTUs) discarding singletons and chimeras in the process. OTUs were compared to the reference sequences of the SILVA 16S database, and taxonomies were predicted considering a minimum confidence threshold of 0.6 using the SINTAX algorithm implemented in USEARCH. Alpha diversity was estimated using the Richness (Observed), Shannon and Simpson indices. Beta diversity was calculated using the Unifrac distance method on the basis of normalized OTU abundance counts per sample. These sample distances were then used in a detrended correspondence analysis (DCA) to reveal possible patterns of inter-sample relations. Alpha and beta diversity calculations and the rarefaction analysis were performed with the R software package phyloseq v1.22.3. To detect differentially abundant OTUs depending on collected sample metadata (e.g., medication, environmental conditions), differential OTU analysis on normalized abundance counts was performed with the R software package DESeq2 v1.18.1. Libraries, sequencing, and data analysis described in this section were performed by Microsynth AG (Balgach, Switzerland).

**RNA-seq analysis**

FACS sorted viable MΦ (300-3000 cells) as well as blood-derived M1 and M2 MΦ (100000 cells) were used for RNA-seq analysis. RNA was isolated using RNeasy micro kit (Qiagen). RNA samples were quantified using Qubit 2.0 Fluorometer (Life Technologies, Carlsbad, CA, USA), and RNA integrity was checked with Agilent TapeStation (Agilent Technologies, Palo Alto, CA, USA). RNA library preparations, sequencing reactions, and initial bioinformatics analysis were conducted at GENEWIZ, LLC. (South Plainfield, NJ, USA). The SMART-Seq v4 Ultra Low input kit for sequencing was used for full-length cDNA synthesis and amplification (Clontech, Mountain View, CA) according to the manufacturer's protocol. Illumina Nextera XT library was used for library preparation. Briefly, cDNA was fragmented, and adaptors were added using transposase, followed by limited-cycle PCR to enrich and add index to the cDNA fragments. Final libraries were analyzed on the Agilent TapeStation for library sizing and quantified using the Qubit dsDNA HS assay kit and by qPCR using the KAPA library quantification kit. The sequencing libraries were multiplexed and clustered on two lanes of a flow cell. After clustering, the flow cell was loaded on the Illumina HiSeq instrument according to the manufacturer's instructions. The samples were sequenced using a 2x150 Paired End (PE) configuration. Image analysis and base calling were conducted by the HiSeq Control Software (HCS). Raw sequence data (.bcl files) generated from Illumina HiSeq were converted into fastq files and de-multiplexed using Illumina's bcl2fastq 2.17 software. One mismatch was allowed for index sequence identification. After investigating the quality of the raw data, sequence reads were trimmed to remove possible adapter sequences and nucleotides with poor quality using Trimmomatic v.0.36. The trimmed reads were mapped to the human reference genome available on ENSEMBL using the STAR aligner v.2.5.2b. The STAR aligner uses a splice aligner that detects splice junctions and incorporates them to help align the entire read sequences. BAM files were generated as a result of this step. Unique gene hit counts were calculated by using featureCounts of the Subread package v.1.5.2. Only unique reads that fell within exon regions were counted. After extraction of gene hit counts, the gene hit counts table was used for downstream differential expression analysis. All statistical analyses were performed using R project software. Comparison of gene expression between the groups of samples was performed with the package DESeq2. The Wald test was used to generate p-values and Log2 fold changes. Genes with adjusted p-values $< 0.05$ and absolute log2 fold changes $> 1$ were regarded as differentially expressed genes for each comparison. The statistically significant set of genes was subjected to a gene ontology analysis by implementing the software GeneSCF. The mgi GO list was used to cluster the set of genes based on their biological process and to determine their statistical significance. A PCA analysis was performed using the plotPCA function of the DESeq2 R package. The plot shows the samples in a 2D plane spanned by their first two principal components. The top 500 genes, selected by highest row variance, were used to generate the plot.

**Statistical analysis**

Data were analyzed using Prism GraphPad 6.0 software and Stata (StataCorp. 2015. Stata Statistical Software: Release 15. College Station, TX: StataCorp LP).

Data are reported as the means ± standard error of the mean (SEM). Statistical significance was determined using unpaired nonparametric two-tailed Mann-Whitney test, and differences were considered statistically significant with * $p \leq 0.05$; ** $p \leq 0.01$; *** $p \leq 0.001$, and **** $p \leq 0.0001$. Unless otherwise noted, figures show pooled patient data from several independent experiments or a representative of repeated experiments. The characteristics and risk factors of study participants were compared between fiber-low versus fiber-high HSCR patient groups, and between subjects with enterocolitis versus no enterocolitis during follow-up, using two-sided Wilcoxon rank-sum test and $\chi 2$-test, as appropriate.

**Data availability**

The datasets generated and analyzed during the current study have been deposited in NCBI Sequence Read Archive under the accession code PRJNA552657 (RNA-seq datasets) and PRJNA550537 (16s rDNA datasets). All other study-related data are available from the corresponding author upon reasonable request.

All authors had access to the study data and reviewed and approved the final manuscript.
